# Reduced synaptic activity and dysregulated extracellular matrix pathways are common phenotypes in midbrain neurons derived from sporadic and mutation-associated Parkinson’s disease patients

**DOI:** 10.1101/2021.12.31.474654

**Authors:** Shani Stern, Shong Lau, Andreea Manole, Idan Rosh, Menahem Percia, Ran Ben Ezer, Maxim N. Shokhirev, Fan Qiu, Simon Schafer, Abed Mansour, Tchelet Stern, Pola Ofer, Yam Stern, Ana Mendes Diniz, Lynne Randolph Moore, Ritu Nayak, Aidan Aicher, Amanda Rhee, Thomas L. Wong, Thao Nguyen, Sara B. Linker, Beate Winner, Beatriz C. Freitas, Eugenia Jones, Cedric Bardy, Alexis Brice, Juergen Winkler, Maria C. Marchetto, Fred H. Gage

## Abstract

Several mutations that cause Parkinson’s disease (PD) have been identified over the past decade. These account for 15-25% of PD cases; the rest of the cases are considered sporadic. Currently, it is accepted that PD is not a single monolithic disease but rather a constellation of diseases with some common phenotypes. While rodent models exist for some of the PD-causing mutations, research on the sporadic forms of PD is lagging due to a lack of cellular models. In our study, we differentiated PD patient-derived dopaminergic (DA) neurons from induced pluripotent stem cells (iPSCs) of several PD-causing mutations as well as from sporadic PD patients. Strikingly, we observed a common neurophysiological phenotype: Neurons derived from PD patients had a severe reduction in the rate of synaptic currents compared to those derived from healthy controls. While the relationship between mutations in genes such as the *SNCA* and *LRRK2* and a reduction in synaptic transmission has been investigated before, here we show evidence that the pathogenesis of the synapses in neurons is a general phenotype in PD. Analysis of RNA sequencing results displayed changes in gene expression in different synaptic mechanisms as well as other affected pathways such as extracellular matrix-related pathways. Some of these dysregulated pathways are common to all PD patients (monogenic or idiopathic). Our data, therefore, shows changes that are central and convergent to PD and suggests a strong involvement of the tetra-partite synapse in PD pathophysiology.

## Introduction

Parkinson’s disease (PD) was first described in 1817 by James Parkinson ^1^, who wrote about patients with shaking palsy, i.e., involuntary tremulous motion with lessened muscular power. PD occurs in approximately 2 of 1,000 people and is highly correlated with aging, affecting about 1% of the older population above 60 years of age ^2, 3^ The main neuropathological symptoms are α-synuclein-containing Lewy bodies and loss of dopaminergic (DA) neurons in the substantia nigra pars compacta. PD patients experience movement difficulties with three cardinal signs: tremor, rigidity and, bradykinesia. Some of the main non-motor symptoms of PD include loss of smell, depression, sleep disorders, and dementia, but a wide range of other symptoms such as excessive saliva ^4^ and susceptibility to melanoma ^5^ may also be present. With the progression of the disease, Lewy body pathology spreads to many areas of the brain, and Lewy bodies containing α-synuclein aggregates are seen throughout many brain areas such as the hippocampus, hypothalamus, neocortex, and cortex ^6–8^.

Approximately 15% of PD cases report familial inheritance of the disease. This percentage may vary significantly in different populations ^9^. Hundreds of variants have been observed in several PD genes that showed a clear association, including *α-synuclein* (*SNCA, PARK4*), *parkin* (*PARK2*), *UCH-L1* (*PARK5*), *PINK1* (*PARK6*), *DJ-1* (*PARK7*), *LRRK2* (*PARK8*), *ATP13A2* (*PARK9*), *GBA, VPS35* (*PARK17*), *EIF4G1*, and *PARK16* ^10, 11^. Thus, PD is not a single disease but a constellation of phenotypes that are displayed variably by patients with different co-morbidities but some commonalities ^12^. Some of the genetically characterized forms of the disease are rapidly evolving, with early-onset ages of the mid-30s or mid-40s ^13–15^, but some genetic forms have a similar age-onset to idiopathic PD ^16^. α-Synuclein immunohistochemistry is currently considered one of the gold standards in the neuropathological evaluation of PD. Aggregates of misfolded α-synuclein in Lewy bodies in DA neurons are now considered a hallmark of PD, but it is not clear whether these Lewy bodies are the cause of neuronal atrophy or a byproduct of the disease ^17–19^. In fact, injection of synthetic α-synuclein fibrils into the dorsal striatum of wild-type mice was enough to elicit and transmit disease pathology and neurodegeneration^20^. Similarly, mice carrying mutations that disrupt physiological tetramers of the α-synuclein protein develop brain pathology and neurodegeneration typical of PD ^21^. In neurons, α-synuclein physiologically localizes mainly to presynaptic terminals ^22–24^. The aggregates of α-synuclein have been observed in cell soma but also in neurites ^25^, and they are widespread in various brain areas in PD patients. However, reports show that micro-aggregates are present in the neurites close to presynaptic terminals, causing synaptic impairment sometimes long before the large aggregates that eventually make up the Lewy bodies form ^17, 26–28^.

Mutations as well as copy number variations in the *SNCA* (*PARK1*) gene, which codes for the α-synuclein protein, have been shown to have a causative effect in PD ^13–15, 29–35^. α-Synuclein is localized mainly in presynaptic terminals ^18^. It helps to maintain the size of the presynaptic vesicular pool as well as vesicle recycling ^36, 37^, and it functions to help neurotransmitter release, especially dopamine ^38–40^. Animal models for mutations and copy number variations in the SNCA gene recapitulate the human motor symptoms and neuronal loss ^41–43^. Surprisingly, there is an almost immediate neurophysiological phenotype of a reduction in synaptic activity after the introduction of different SNCA mutations in rodent models ^37, 44–48^. A reduction in synaptic activity has been shown on its own to cause neuronal atrophy in different types of neurons ^37, 48, 49^, suggesting a positive feedback mechanism that further increases neuronal cell death.

Leucine-rich repeat kinase 2 (*LRRK2, PARK8*) is another gene with a causative association to PD ^10^. It is the most common form of genetic PD, and mutations are highly prevalent in certain populations^50^. The precise physiological function of *LRRK2* is not completely understood, but recent studies have shown that *LRRK2* is involved in cellular functions such as neurite outgrowth, cytoskeletal maintenance, vesicle trafficking, autophagic protein degradation, and the immune system ^51^. Drosophila models with overexpression of *LRRK2* recapitulate PD phenotypes of motor dysfunction and DA cell death ^52, 53^ and so do C. elegans overexpression models ^54^. However, flies and nematodes do not express a-synuclein and are therefore not a great model for studying PD. Rodent models with *LRRK2* mutations show minimal evidence of neurodegeneration ^55, 56^ but do show locomotor impairments ^56^ and reductions in stimulated dopamine neurotransmission and D2 receptor function. Interestingly, *LRRK2* was found to regulate synaptic vesicle endocytosis and recycling and neurotransmitter release ^57–59^. RNA-mediated silencing of *LRRK2* affected postsynaptic currents as well as presynaptic vesicle trafficking and recycling ^60^.

Parkin (encoded by *PARK2*) is a ubiquitin-ligase enzyme expressed in the CNS and peripheral tissues ^61^. It is a multifunctional protein that is involved in many intracellular processes, and several substrates for it have been identified. Parkin is localized on synaptic vesicles and displays a distribution pattern similar to that of synapsin I, a protein that associates with the cytoplasmic surface of synaptic vesicles ^62, 63^. Differential fractioning of rat brain lysates revealed that parkin was enriched in the fraction containing PSD-95, a postsynaptic marker ^64^. Homozygous and heterozygous mutations are considered risk factors for early-onset PD, though the contribution of the heterozygous form to the onset of PD is still considered controversial ^65–67^. Loss of function mutations in the *PARK2* gene cause early and severe degeneration of DA neurons of the substantia nigra pars compacta ^68^. Parkin has been shown to negatively regulate the number and strength of excitatory synapses in rats ^69^ and to ubiquitinate synaptic proteins ^70^. Pathogenic parkin mutations were shown to disrupt glutamatergic synaptic transmission and plasticity by impeding NMDA and AMPA receptor trafficking ^71, 72^. Mutations in Parkin may therefore cause dysregulation of glutamatergic and DA synapses that eventually culminates in DA neuron cell death.

Overall, quite a few animal model studies suggest that several PD-associated mutations result in a synaptic pathology that occurs before neuronal cell death and may even be the cause of neuronal degeneration ^27^. These studies are further supported by other neuroanatomical studies of post-mortem patient brain samples from familial PD cases ^73–75^. Here, we used PD patient-derived neurons to study the neuropathology in PD using the induced pluripotent stem cells (iPSCs) technique. Reprogramming adult cells into stem cells is known to erase aging signatures ^76, 77^, and therefore the neurons derived in this method can be considered as young, and even pre-natal, neurons. We found that in all the PD lines (both mutations-derived and sporadic PD (sPD)) there is a significant reduction in the rate of synaptic events in these “young” neurons. The use of iPSCs allowed us to also measure neurons derived from PD patients with no known genetic mutations, who are therefore considered sporadic. Our cohort consisted of a few patients with sPD. A few of the patients had mutations in the SNCA gene (one patient with a duplication, one patient with a triplication, and a genetically engineered line with the A53T mutation). One patient had a mutation in the LRRK2 gene, and two patients had a mutation in their Parkin gene. Our results suggest a biological predisposition that exists in PD patients, even without other epigenetic or environmental influences. Furthermore, our methods allowed us to recognize gene expression patterns and gene ontology (GO) terms that are common to neurons derived from patients with different PD mutations as well as to sPD.

## Results

### A drastic reduction in synaptic activity is observed in neurons derived from patients with duplication and triplication of the SNCA gene

Our first cohort consisted of a control line (healthy individual, the 40102 line), a PD patient with a duplication of the α-synuclein gene (denoted as 2X), and a PD patient with a triplication of the α-synuclein gene (denoted as 3X). The patient with the α-synuclein triplication had early-onset, autosomal dominant PD at age 38 and was previously described ^91^. We differentiated these lines into DA neurons (see Methods) and used whole-cell patch-clamp to assess intrinsic properties as well as synaptic activity in the neurons. An immunostaining image for DAPI to mark cell bodies, MAP2 to mark neurons, and TH to specifically mark dopaminergic neurons is shown in Figure 1Q. A total of 15 control neurons, 11 2X neurons, and 10 3X neurons were patch clamped approximately 50 days after the start of differentiation. The total number of evoked potentials in the 17 first depolarization steps was decreased in the 2X neurons counting (see Methods, Total evoked action potentials) (p=0.013, Fig. 1A). Representative traces of evoked action potentials are shown in Figure 1B-1D. Next, we used voltage-clamp mode to measure the sodium and potassium currents in the control and PD neurons; representative traces of the currents are shown in Figure 1E-1G. The average sodium currents are presented in Figure 1H. At −30 mV, control neurons displayed a sodium current of 1.5±0.6 pA/pF, whereas 2X neurons displayed a sodium current of 1±0.07 pA/pF, and 3X neurons displayed a sodium current of 13.2±6 pA/pF (p=0.026 compared to controls). At - 20 mV, control neurons displayed a sodium current of 16.2±3 pA/pF, whereas 2X neurons displayed a sodium current of 4.2±1.8 pA/pF (p=0.006 compared to controls), indicating the opening of the sodium channels at voltages between control and the PD neurons. The slow potassium currents are presented in Figure 1I. An ANOVA indicated no significant differences. The fast potassium currents are shown in Figure 1J. Similarly, running an ANOVA did not indicate significant differences. Analysis of the shape of the action potential is presented in Supplementary Figure 1A-D.

**Figure 1.**
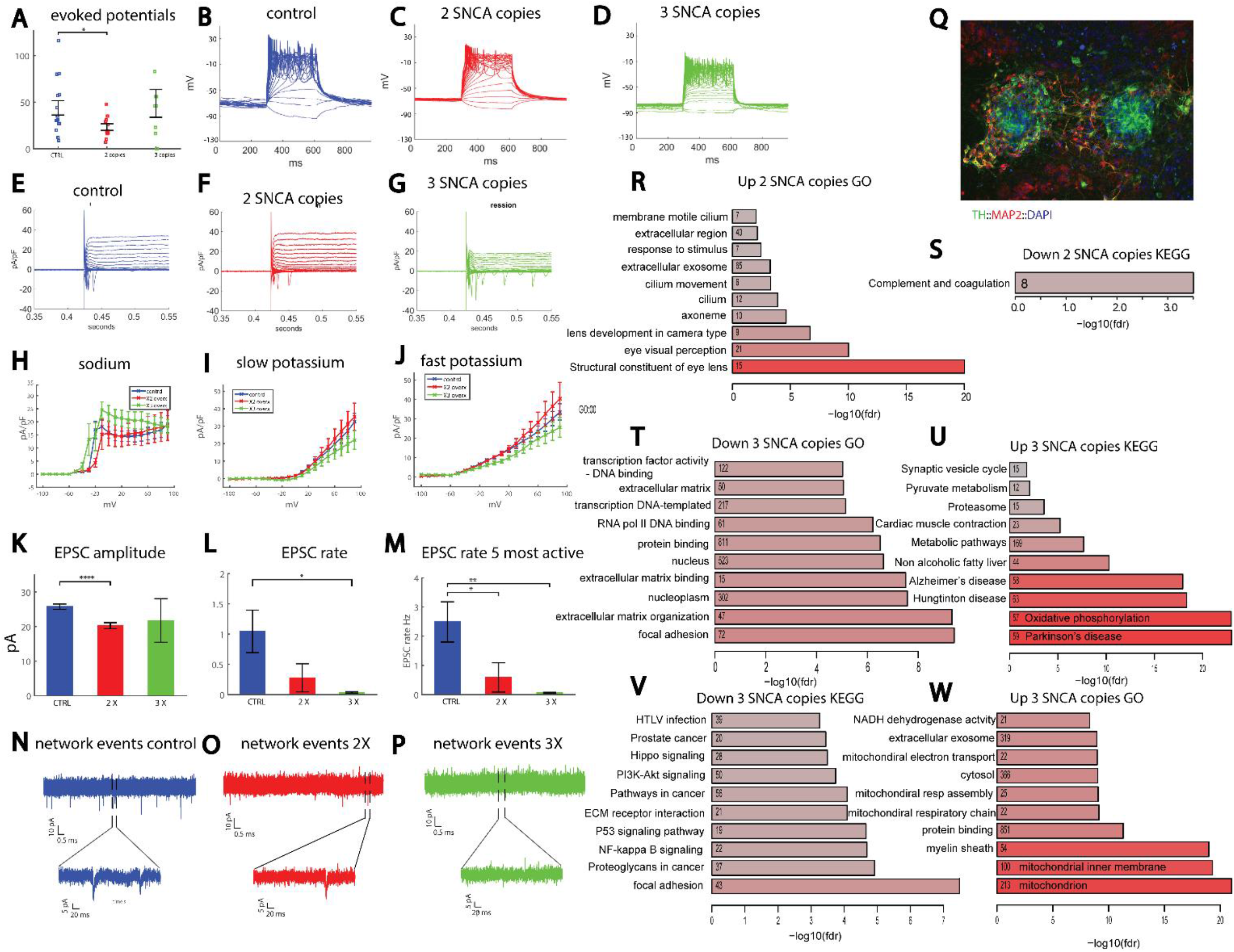
Reduced synaptic activity in neurons of patients with two or three copies of the SNCA gene (denoted as 2X and 3X). A. Neurons derived from the 2X patient display reduced excitability compared to controls. B-D. Representative evoked potential recordings of a control (B), 2X patient (C), and 3X patient (D). E-G. Representative traces of sodium and potassium currents in control individuals (E), 2X patient (F), and 3X patient (G). H. Sodium currents of neurons from 3X patient opened at a less depolarized potential than control DA neurons, whereas those derived from the patient with the double copy opened at a more depolarized potential. I. Slow potassium currents are reduced in DA neurons derived from 3X patient compared to controls. J. Fast potassium currents are reduced in DA neurons derived from 3X patient compared to controls. K. The mean amplitude of synaptic currents is reduced in DA neurons from 2X patient compared to controls. L. The average rate of synaptic events is reduced in DA neurons derived from 2X patient and further reduced in 3X patient. M. The differences in the EPSC rate become significant when we analyze the neurons with the highest rate of synaptic currents, even in the 2X patient. This arises from the fact that diversity in the rate of synaptic currents is very large in neuronal cultures, since more than 50% of the neurons have very few synaptic currents, and only around 30% of the neurons are highly connected, driving the network activity. N-P. Example traces of recordings of synaptic currents in DA neurons derived from a control individual (N), the 2X patient (O), and the 3X patient (P). The lower image in each is a zoom of the segment denoted by the black bars in the upper image. Q. An example image of a control culture stained for TH (marking DA neurons), MAP2 (marking all neurons), and DAPI (marking all cells). R-W. Bar plots showing top overrepresented terms or pathways among differential gene sets. Numbers indicate counts of overlapping genes; red color represents the significance of the overlap. R. Up-regulated GO terms sets in the 2X patient compared to controls. S. Down-regulated KEGG gene set in the 2X patient compared to controls. T. Down-regulated GO terms in the 3X patient compared to controls. U. Up-regulated KEGG pathways in the 3X patient compared to controls. V. Down-regulated KEGG pathways in the 3X patient compared to controls. W. Up-regulated GO terms in the 3X patient compared to controls. In this figure and the next figures, asterisks represent statistical significance by the following code: * p value<0.05, **p value<0.01, ***p<0.001, ****p<0.0001. Error bars represent the standard error in this figure.

Recording of EPSCs revealed an altered network activity in both the 2X and 3X neurons. The cumulative distribution of the amplitudes of EPSCs was left-shifted in 2X neurons and further left-shifted in 3X neurons, indicating lower amplitudes of EPSCs (Supplementary Fig. 1E). The average EPSC amplitude was decreased in the 2X neurons (p=0.0025 between control and 2X neurons, Fig. 1K). The rate of synaptic events per recorded cell is shown in Figure 1L. The average rate of EPSCs was 1.04±0.35 Hz for control neurons, 0.28±0.23 Hz for 2X neurons, and 0.037±0.012 Hz for 3X neurons. Although the differences may seem large, the diversity in the rate of EPSCs varied greatly within neurons; therefore, a significant change was only seen in the 3X neurons (p=0.03 compared to controls, Fig. 1L, and p=0.1 for 2X neurons compared to controls). To decrease the diversity, we also calculated the average rate of EPSCs, when only the five most active neurons within a group were taken into the calculation. Here, the averages were 2.5±0.7 Hz for control neurons, 0.6±0.5 Hz for 2X neurons, and 0.06±0.02 Hz for 3X neurons (p=0.05 for 2X compared to control neurons, and p=0.007 for 3X compared to control neurons, Fig. 1M). Figure 1N displays an example recording of EPSCs in a control neuron, Figure 1O displays an example recording of EPSCs in a 2X neuron, and Figure 1P displays an example recording of a 3X neuron. Cell capacitance was not significantly different between the 3 groups (Supplementary Fig. 1G). The input conductance was increased in 3X neurons (p=0.0016 between 3X and control neurons, and p=0.0019 between 2X and 3X neurons, Supplementary Fig. 1H). To summarize, a reduction in synaptic activity was observed and exaggerated with α-synuclein gene dosage.

Analyzing the RNA sequencing data and looking at gene ontology and affected KEGG and MsigDB pathways, we found quite a few dysregulated pathways. Four control lines were pooled together for this analysis; 40102, UKERf1JF-X-001, UKERf33Q-X-001, and UKERfO3H-X-001. We plotted the 15 most significant pathways in Figures 1R – 1W. The differentially regulated pathways are denoted to the left, with the number of genes that are involved inside the bars. The full lists of GO terms, KEGGS, and MSigDB pathways are presented in Supplementary Figures 2-4. Interestingly, in the patient with the SNCA triplication, there were many synapse-related dysregulated pathways, with dozens of affected genes (Fig. 1U and Supplementary Fig. 2, 4), such as synapse part, synaptic vesicle cycle, synaptic transmission, and many more. In the neurons derived from this patient, we also saw a very severe reduction in the rate of synaptic currents, further confirming that synapses were severely disrupted. The full list of the genes involved in the affected pathways is presented in Supplementary Figure 3 and consists of both presynaptic and postsynaptic genes. We hypothesize that PD neurons are unable to form or maintain effective synapses, which we detect as a reduction in the rate of synaptic activity with electrophysiology, and this failure, in turn, triggers compensation mechanisms that cause further dysregulation of synapse-related genes. Additionally, in the upregulated pathways, we found “Parkinson’s disease,” Alzheimer’s disease” and “Huntington disease” showing commonalities between these neurodegenerative diseases. Several oxidative phosphorylation-related pathways were also significantly overlapped with upregulated genes, supporting the known role of oxidative stress in PD, as well as mitochondrial genes that were dysregulated. In the pathways significantly overlapped with downregulated genes, we found several metabolic pathways and pathways related to protein folding, as well as a few pathways that were not known before to relate to PD and are common in many of our mutation lines as well as sPD (as will be presented in the subsequent figures), such as extracellular matrix pathways (probably the most dysregulated pathways), and PI3K-Akt signaling pathways, lysosome membrane degradation, and focal adhesion.

In the neurons derived from the patient with the SNCA duplication, the “ion transport” pathway and other transporter pathways were dysregulated (See Supplementary 2,4), which may be related to the changes that we observed in neuronal excitability and ionic currents. In the electrophysiology data, the reduction in synaptic activity was less pronounced than in the patient with the SNCA triplication, further emphasizing the need for electrophysiology for targeting smaller changes that are related to functionality. Several extracellular matrix-related pathways were also dysregulated in the patient with the SNCA duplication, further suggesting that there might be changes to the extracellular matrix structure that complement or even precede a possible synaptic degradation. Overall dysregulation of extracellular matrix pathways is common to all the PD lines that we worked with.

### A drastic reduction in synaptic activity is observed in DA neurons derived from patients with LRRK2 and Parkin mutations

Our second cohort consisted of two control (two healthy subjects, 40102 and UKERfO3H-X-001 lines) individuals, one patient with a mutation in the LRRK2 and two patients with mutations in the Parkin gene. We differentiated these lines using a DA human neuronal protocol ^78^ (see Methods) and used whole-cell patch-clamp to assess intrinsic properties as well as synaptic activity in the neurons (n=20 control neurons, n=17 from the patient with the LRRK2 mutation, n=12 from one patient with a Parkin mutation and n=15 neurons from the second patient with the Parkin mutations). The total number of evoked potentials in the 17 first depolarization steps was decreased in the LRRK2 DA neurons (see Methods, Total evoked action potentials) and in the neurons with the Parkin mutation. (p=0.0012 between control and LRRK2 mutation, p=0.06 between control and the two Parkin mutations, pooling the data of the two mutations together, Fig. 2A). Representative traces of evoked action potentials are shown in Figure 2B-2E. Next, we measured the sodium and potassium currents in voltage-clamp mode in the control, LRRK2, and Parkin DA neurons. The average sodium currents are presented in Figure 2F. The average slow potassium currents are presented in Figure 2G and the average fast potassium currents are presented in Figure 2H. Representative traces of measured EPSCs are shown in Figure 2I (control), 2J (LRRK2 mutation), 2K (first patient with a Parkin mutation termed Parkin1), and 2L (second patient with a Parkin mutation, termed Parkin2). There was a significant reduction in the rate of synaptic currents in all three PD-causing mutations. Control neurons had 0.95±0.25 Hz, LRRK2 0.3±0.16 Hz, Parkin1 0.16±0.07 and Parkin2 0.21±0.08 Hz (LRRK2 compared to control p=0.05, Parkin1 compared to control p=0.027, Parkin2 compared to control p=0.019, Fig. 2M). There was no significant change in the average amplitude of the synaptic currents (Fig. 2N).

**Figure 2.**
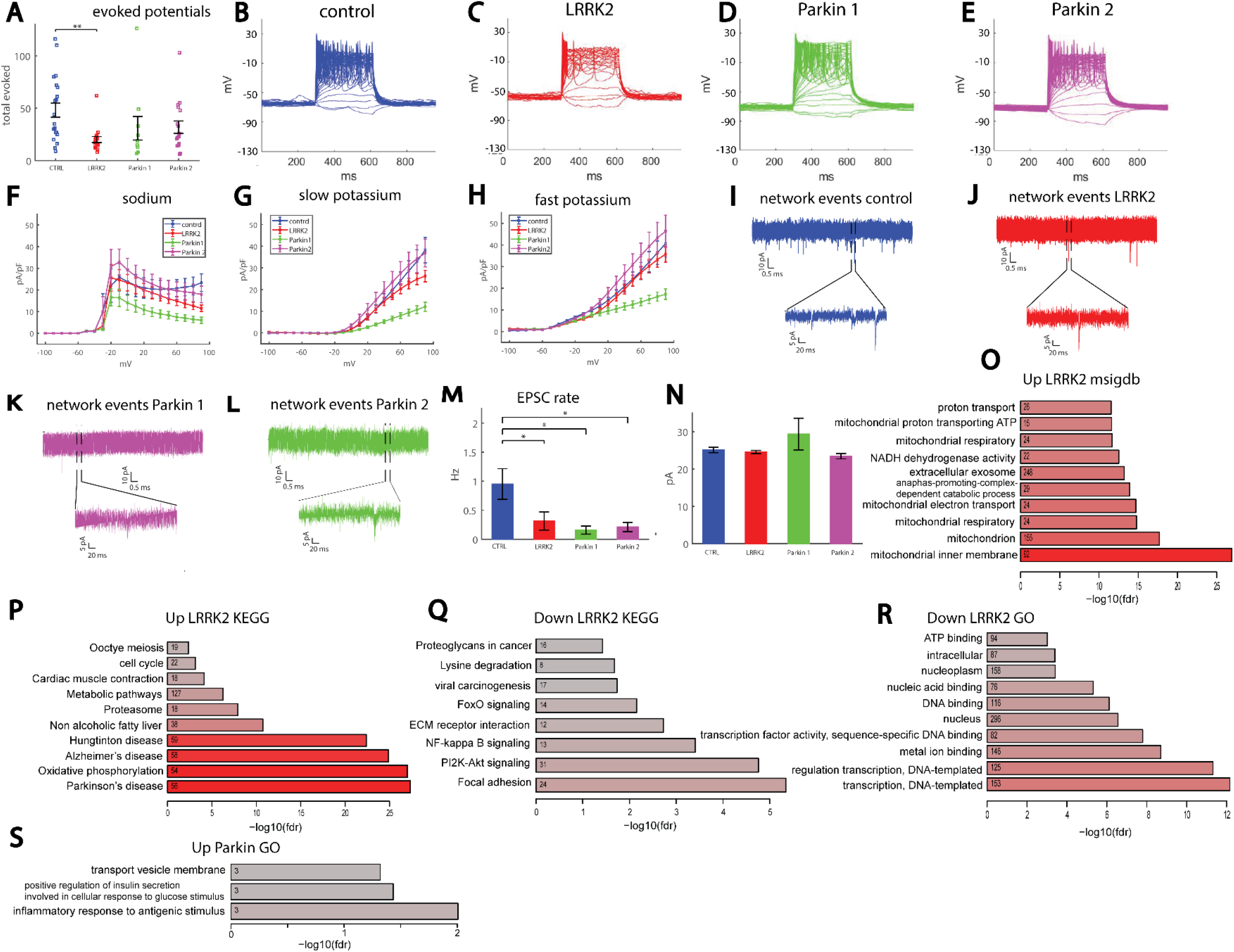
Reduced synaptic activity in neurons derived from patients with LRRK2 and Parkin mutations compared to control dopaminergic neurons. A. Neurons derived from a patient with a LRRK2 mutation (CHE line) display reduced excitability measured by the total evoked potentials compared to controls. The 2 patients with the Parkin mutations had a reduction in excitability that was not statistically significant (Parkin1 and Parkin2 patient lines). B-E. Representative traces of recordings of evoked potentials in a control neuron (b), LRRK2 neuron (c), Parkin1 neuron (d), and Parkin2 neuron (e). f. Sodium currents were reduced in one of the patients with the Parkin mutations (Parkin1) but not in the other patient with Parkin and LRRK2 mutations compared to controls. G. Slow potassium currents were significantly reduced in neurons derived from the first patient with a Parkin mutation (Parkin1) but not in the second patient with a Parkin mutation (Parkin2) and LRRK2 mutations compared to controls. H. Fast potassium currents were significantly reduced in neurons derived from the patient with the first Parkin mutation (Parkin1) but not in the second patient with a Parkin mutation (Parkin2) and LRRK2 mutations compared to controls. I. Example recording of synaptic currents in a neuron derived from a healthy control. J. Example recording of synaptic currents in a neuron derived from a patient with a LRRK2 mutation (lower plot is a zoom in on the segment denoted in the black segmented lines in the upper graph). K. Example recording of synaptic currents in a neuron derived from the first patient with a Parkin mutation (Parkin1, ower plot is a zoom in on the segment denoted in the black segmented lines in the upper graph). L. Example recording of synaptic currents in a neuron derived from the second patient with a Parkin mutation (Parkin2, lower plot is a zoom in on the segment denoted in the black segmented lines in the upper graph). M. The average rate of synaptic events is reduced in neurons derived from a patient with a mutation in the LRRK2 gene and further reduced in patients with a mutation in the Parkin gene. N. No significant change is observed in the mean amplitude of the neurons with the LRRK2 and Parkin mutations compared to control. o-t. Bar plots showing top overrepresented terms or pathways among differential gene sets. Numbers indicate counts of overlapping genes; red color represents the significance of the overlap. O. Up-regulated GO terms in the neurons of the LRRK2 patient compared to controls. P. Upregulated KEGG pathways in neurons of LRRK2 patient compared to controls. Q. Down-regulated KEGG pathways in neurons of LRRK2 patient compared to controls. R. Down-regulated GO terms in the DA neurons of the Parkin patient compared to controls. S. Up-regulated GO terms in the DA neurons of the Parkin patient compared to controls. In this figure asterisks represent statistical significance by the following code: * p value<0.05, **p value<0.01. Bars represent the standard error in this figure.

Additionally, we analyzed several features of the shape of the spikes. There was a significant decrease in the spike amplitude in the LRRK2 mutation (LRRK2 vs. control, p=0.01; Parkin pooled together vs. control, p=0.07, Supplementary Fig. 5A). There were no significant changes in the spike threshold (Supplementary Fig. 5B), in the spike width (Supplementary Fig. 5C), or the fast AHP amplitude (Supplementary Fig. 5D). The cumulative distribution of the amplitude of EPSCs was similar between the control and the LRRK2 and Parkin lines (Supplementary Fig. 5E). There were no significant differences in the capacitance of the neurons (Supplementary Fig. 5F), and the input conductance was significantly larger in the first line with the Parkin mutation compared to the control (p=0.03, Supplementary Fig. 5F).

The top 10 GO terms and KEGG and MSigDB over-representation analyses for the DA neurons with the LRRK2 and Parkin mutations are presented in Figure 2O-S, and the full list is shown in Supplementary Figures 2,4. The differentially expressed gene list is presented in Supplementary Figure 3. Interestingly, several synaptic transmission pathways were significantly overlapped with upregulated genes in the LRRK2 mutation, in agreement with our electrophysiological data, with approximately 10-20 affected genes in each of these pathways; these include “regulation of synaptic transmission”, “excitatory synapse” and more. Similar to the neurons derived from patients with the SNCA copy number variation, Parkinson’s disease, Alzheimer’s disease, and Huntington’s disease, oxidative phosphorylation, and oxidation reduction-related pathways were significantly overrepresented among upregulated genes. In addition, mitochondrial pathways were overrepresented among upregulated genes, presenting more evidence that mitochondrial defects are present in PD. Among the terms significantly overlapping with downregulated genes we found many affected pathways that were related to the extracellular matrix, PI3K-Akt signaling pathway, lysine degradation, focal adhesion, and FoxO signaling pathways, as well as protein folding-related pathways. Some of these pathways were not previously known to be associated with PD, but they were misregulated in most of our PD lines, showing new possible cellular defects that need to be targeted for treatment. For the DA neurons derived from the PD patient with the Parkin mutation, there were not many significantly dysregulated pathways, but these did include “transport vesicle membrane.”

### A reduction in synaptic activity is observed in neurons derived from sPD patients

We continued the study by differentiating neurons from a PD patient with no PD-causing mutations (UKERfAY6-X-001). We recorded from n=20 control neurons (two patients: 40102 and UKERfO3H-X-001) and n=22 neurons from the sPD patient. There was no significant change in the total evoked action potentials (Fig. 3A for averages and 3B, 3C for representative traces). The average of the sodium currents is presented in Figure 3D. The average over the slow potassium currents is presented in Figure 3E and the average of the fast potassium currents is presented in Figure 3F. Representative EPSC traces are presented in Figures 3G and 3H. There were no significant differences in the mean amplitude (Fig. 3I) or the mean rate (Fig. 3J) of EPSCs. However, when counting the number of neurons that had synaptic activity (see methods), we did see a significant reduction in this number in the sPD neurons (p=0.04, Fig. 3K). Furthermore, analysis of the decay time constant of the synaptic events reveals a faster decay in sPD neurons with an average of 2.3±0.1 ms for control neurons and 1.5±0.07 ms for sPD neurons (p=9e-12, the distribution is presented in Supplementary Fig. 6). Spike shape analysis did not reveal any significant changes in the spike height (Supplementary Fig. 7A). The spike was significantly narrower in the sPD neurons (control 6±0.8 ms, sPD 3.6±0.2 ms, p=0.0035, Supplementary Fig. 7B). The spike threshold was significantly more negative in the sPD neurons (control −18.4±1.3 mV, sPD −24.6±1 mV, p=0.0006, Supplementary Fig. 7C). There was no significant change in the fast AHP (Supplementary Fig. 7D) or the capacitance (Supplementary Fig. 7F), and the input conductance was larger in sPD neurons, but not significantly (control 0.45±0.12nS, sPD 0.87±0.17 nS, p=0.06, Supplementary Fig. 7G). The cumulative distribution of amplitudes of EPSCs was similar between the control and sPD neurons (Supplementary Fig. 7E).

**Figure 3.**
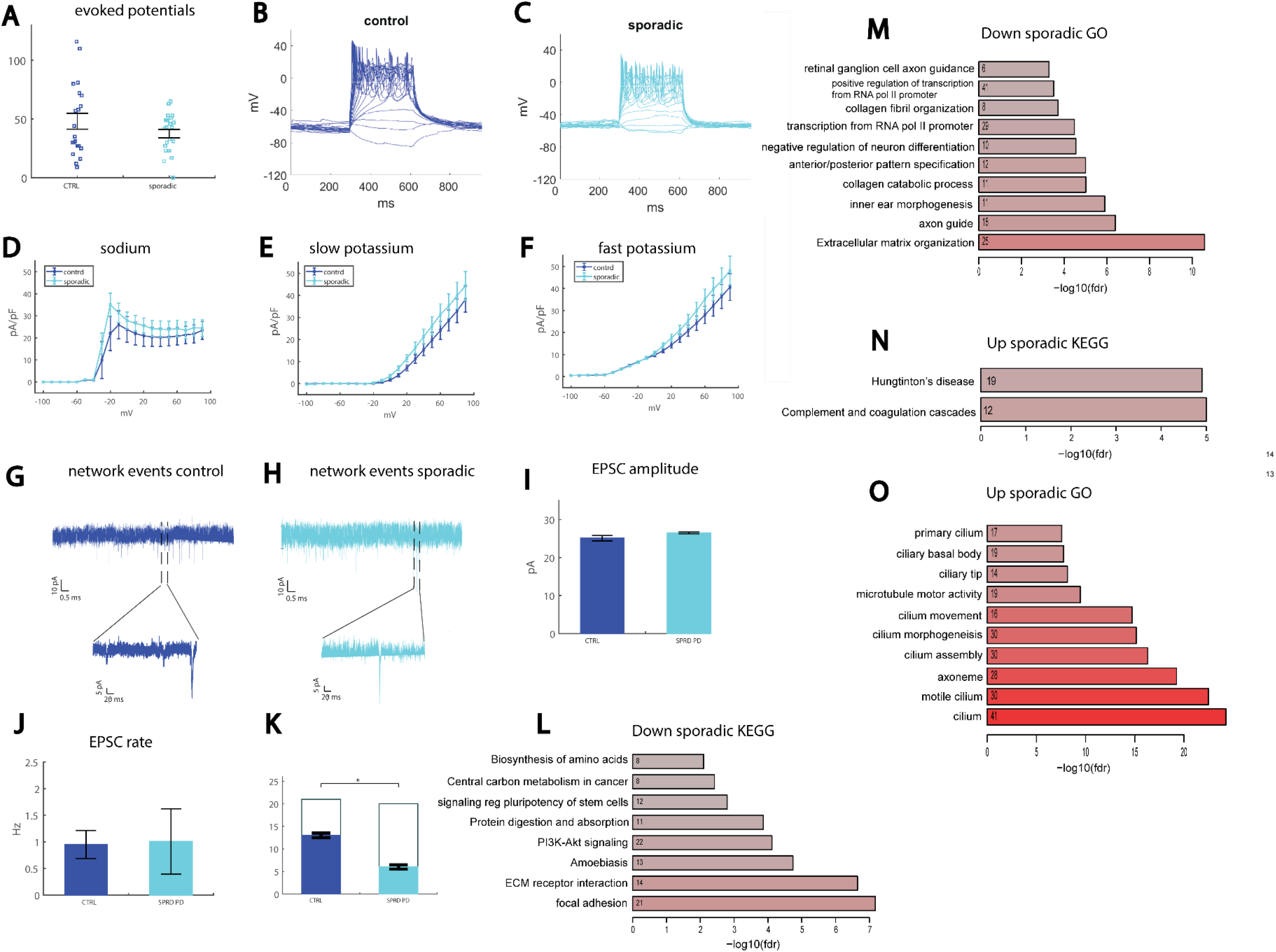
A reduction in the number of neurons with synaptic activity is observed in neurons derived from a sPD patient. A. No significant change was observed in the excitability measured by the total evoked potentials in the neurons derived from the sPD patient compared to controls. B. Representative example of evoked action potentials in a control neuron. C. Representative example of evoked action potentials in a neuron derived from a sPD patient. D. No significant changes were observed in the sodium currents. E. No significant changes were observed in the slow potassium currents. F. No significant changes were observed in the fast potassium currents. G. Example recording of synaptic currents in a neuron derived from a healthy control (the lower plot presents a zoom in on the segment denoted in the black dashed lines in the upper graph). H. Example recording of synaptic currents in a neuron derived from a patient with sPD (the lower plot presents a zoom in on the segment denoted in the black dashed lines in the upper graph). I. No significant change was observed in the mean amplitude of the synaptic currents in the sPD neurons. J. No significant change was observed in the mean rate of the synaptic currents in the sPD neurons. K. A significant reduction in the number of neurons that had synaptic activity was observed in sPD neurons compared to controls. L-O. Bar plots showing top overrepresented terms or pathways among differential gene sets. Numbers indicate counts of overlapping genes; red color represents the significance of the overlap. L. Down-regulated KEGG pathways in neurons of sPD patients compared to controls. M. Down-regulated GO terms in the neurons of the sPD patients compared to controls. N. Up-regulated KEGG pathways in neurons of sPD patients compared to controls. O. Up-regulated GO terms in the neurons of the sPD patients compared to controls. * p value<0.05. Bars represent the standard error in this figure.

Three sPD lines (UKERfRJO-X-001, UKERfAY6-X-001, and UKERfM89-X-001) were pooled together for the analysis of gene expression, plus 4 control (lines 40102, UKERf1JF-X-001, UKERf33Q-X-001, and UKERfO3H-X-001). The top 10 GO terms, KEGG and MSigDB dysregulated pathways for the DA neurons derived from sPD patients are presented in Figure 3L-O, and the full list is shown in Supplementary Figures 2-4. There were a few dysregulated pathways that were related to the synapse such as regulation of synaptic structure, regulation of synaptic organization, regulation of synaptic plasticity, and more. The most dysregulated pathways were related to nasopharyngeal carcinoma, with almost 250 affected genes and an FDR of 1.6e-105. Other highly dysregulated pathways were related to the cilium. Like the other genetic PD lines, the extracellular matrix was highly affected, and more dysregulated pathways that were repeatedly affected in our PD lines were focal adhesion, collagen processes, PI3K-Akt, protein digestion and absorption, and pathways related to reactive oxygen species and metabolic processes. Several hypoxia-related pathways were affected as well.

### A reduction in synaptic activity is observed in neurons derived from more sPD patients using a different protocol and selecting a subset of the neurons

We next analyzed data that were acquired using a different differentiation protocol (see Methods, second DA differentiation); in addition, only neurons that were defined as type 5 neurons (see definition in Methods) were used for the analysis. Strikingly, despite the different methods used both in differentiating the neurons and in analyzing the data, the main phenotype of a reduction in synaptic activity was present in the sPD neurons as well.

We recorded from one control individual (line 40102) and two sPD patients (UKERfRJO-X-001, UKERfR66-X-001). It should be noted that one of the patients had a heterozygous missense mutation in the EIF4G1 gene (UKERfRJO-X-001). The number of total evoked action potentials was not different between control and sPD neurons (averages presented in Fig. 4A and representative traces are presented in Fig. 4B and 4C). The averages of the sodium currents are presented in Figure 4D, the averages of the slow potassium currents are presented in Figure 4E, and the averages of the fast potassium currents are presented in Figure 4F. Representative traces of synaptic activity are presented in Figure 4G (control) and 4H (sPD). The average amplitude of the synaptic currents was increased in sPD neurons, but not significantly (Fig. 4I). The average rate of synaptic currents was significantly reduced in the sPD neurons (control 3±0.6 Hz, sPD 1.4±0.2 Hz, p=0.0037, Fig. 4J). The cumulative distribution of the amplitude of synaptic currents was slightly right-shifted in the sPD neurons. The capacitance was reduced in the sPD neurons, indicating smaller cells (control 64±38 pF, sPD 45±33 pF, p=0.02). Performing action potential shape analysis, we did not observe any significant changes in the spike height (Supplementary Fig. 8A) or the spike width (Supplementary Fig. 8B). The amplitude of the fast AHP was larger in the sPD neurons (control −5.7±0.8 mV, sPD −10.9±0.6 mV, p=0.0000035, Supplementary Fig. 8C), and the threshold for evoking an action potential was more depolarized (control −43±0.8 mV, sPD −39.9±0.6 mV, p=0.0016, Supplementary Fig. 8D).

**Figure 4.**
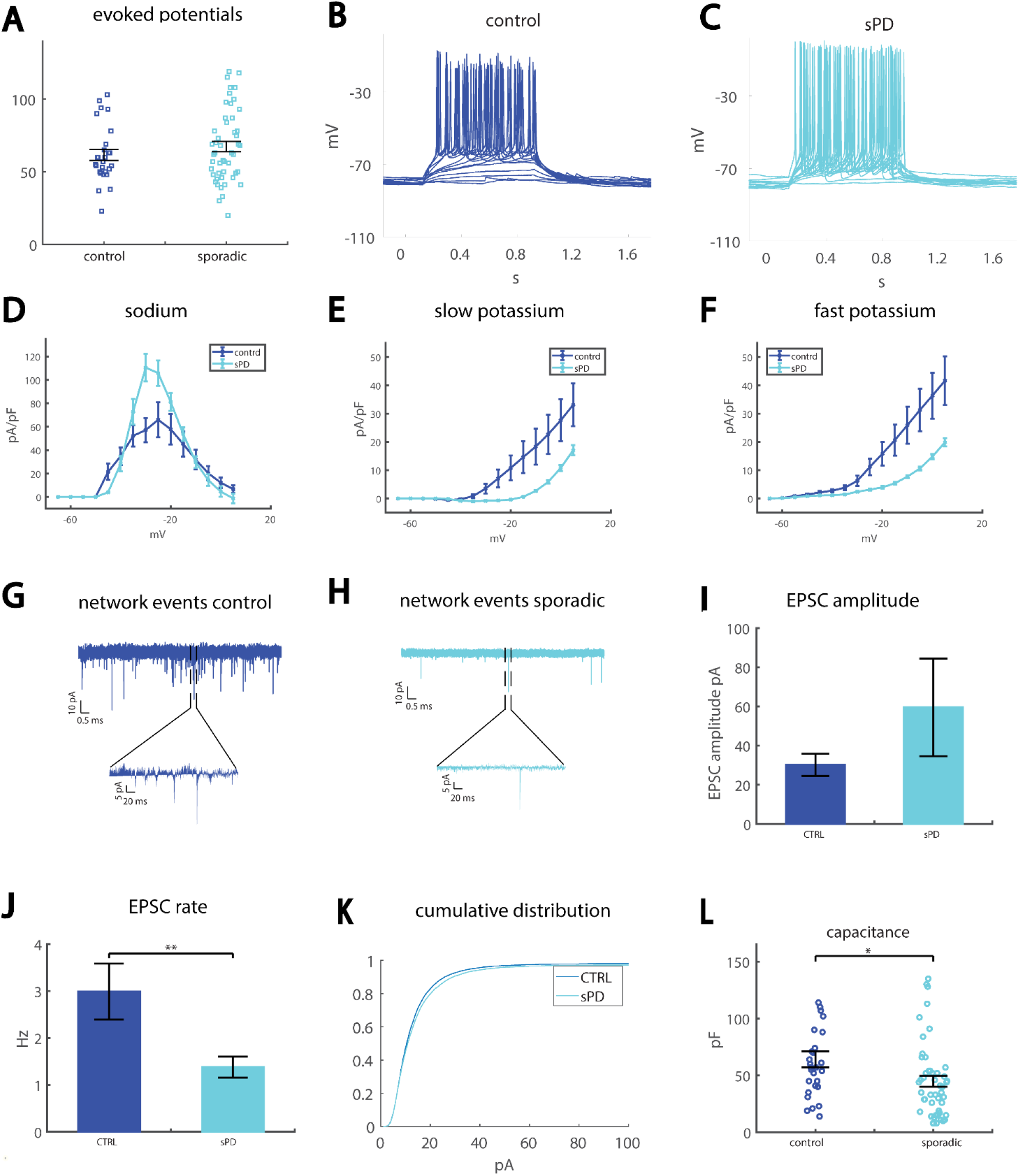
Reduced synaptic activity in neurons derived from 2 sPD patients compared to control neurons. A. No significant change was observed in the excitability measured by the total evoked potentials in the neurons derived from the sPD patient compared to controls. B. Representative example of evoked action potentials in a control neuron. C. Representative example of evoked action potentials in a neuron derived from a sPD patient. D. Sodium currents are increased in the sPD neurons compared to the controls. E. The slow potassium currents are reduced in the sPD neurons compared to the control neurons. F. The fast potassium currents are reduced in the sPD neurons compared to the control neurons. G. A representative trace of synaptic currents in a control neuron (the lower plot presents a zoom-in on the segment denoted in the black dashed lines in the upper graph). H. A representative trace of synaptic currents in an sPD neuron (the lower plot presents a zoom in on the segment denoted in the black dashed lines in the upper graph). I. The mean amplitude of the synaptic currents is increased, but not significantly, in the sPD neurons. J. The mean rate of synaptic currents is significantly reduced in the sPD neurons. K. The cumulative distribution of the sPD neurons is slightly right-shifted, indicating large amplitudes of synaptic currents. L. The capacitance of sPD neurons is significantly reduced in sPD neurons compared to control neurons. * p value<0.05; ** p value<0.01. Bars represent the standard error in this figure.

### A reduction in synaptic activity is observed in neurons derived from an edited iPSC line with an inserted A53T mutation in the SNCA gene

We next performed experiments on neurons derived from a healthy human subject whose fibroblasts were reprogrammed into iPSCs and on an engineered line, in which the A53T mutation was edited into the SNCA gene (isogenic lines). The healthy line and the edited mutated line were differentiated into DA neurons by a differentiation technique that is proprietary to Fujifilm Cellular Dynamics Inc (CDI) (iCell DopaNeurons ^92^). Having an edited healthy line allowed us to observe a similar genetic background, thereby measuring the neuronal changes that occurred specifically due to the A53T mutation. iCell DopaNeurons (CDI) were previously shown to have a protein expression pattern that supported a midbrain lineage DA phenotype ^92^. Using whole-cell patch-clamp, we measured the functional features of A53T neurons compared to the control neurons. Fourteen A53T and 17 control neurons were recorded. The total number of evoked potentials in 17 depolarization steps (see Methods) was not significantly different between A53T neurons (39±5) and control neurons (43±6) (Fig. 5A, and representative traces in Fig. 5B and 5C). Using voltage clamping we measured the sodium/potassium currents in the neurons (representative traces are shown in Fig. 5D and 5E). Sodium currents were not significantly different, except for the opening of the sodium channels; at −30 mV, control neurons displayed a sodium current of 1.6±0.7 pA/pF, whereas A53T neurons displayed a sodium current of 10.4±2.5 pA/pF (p=0.0012, Fig. 5F). Slow and fast potassium currents were not significantly different between A53T neurons and controls (Fig. 5G, 5H). Spike parameters were not altered in the A53T neurons (Supplementary Fig. 9A-D). The cumulative distribution curve of the amplitude of synaptic currents was left-shifted in A53T compared to controls, indicating lower amplitudes of synaptic currents (Fig. 5I). The mean amplitude of synaptic currents was significantly reduced in A53T neurons (16.7±1.2 pA) compared to control neurons (20.6±0.3 pA, p=2e-6, Fig. 5J). The mean rate of synaptic events was significantly reduced in A53T neurons (0.2±0.06 Hz) compared to control neurons (1±0.4 Hz, Fig. 5K, p=0.05). Representative example recordings of synaptic currents for control neurons (Fig. 5L) and an A53T neuron (Fig. 5M) are shown. Cell capacitance was smaller, but not significantly, in A53T neurons (20.2±1.6 pF) compared to control neurons (23±1.4 pF, Supplementary Fig. 9E). Further imaging of neurons stained for TH shows that the A53T neurons were smaller and had fewer, less arborized neurites (Supplementary Fig. 10A-H). The input conductance was decreased, but not significantly, in A53T neurons (Supplementary Fig. 9F). The mean percentage of neurons with neurite beading was 3.7±0.6 % in control cultures and 28±1.8 % in A53T cultures (p=2e-10, Supplementary Fig. 10I-K).

**Figure 5.**
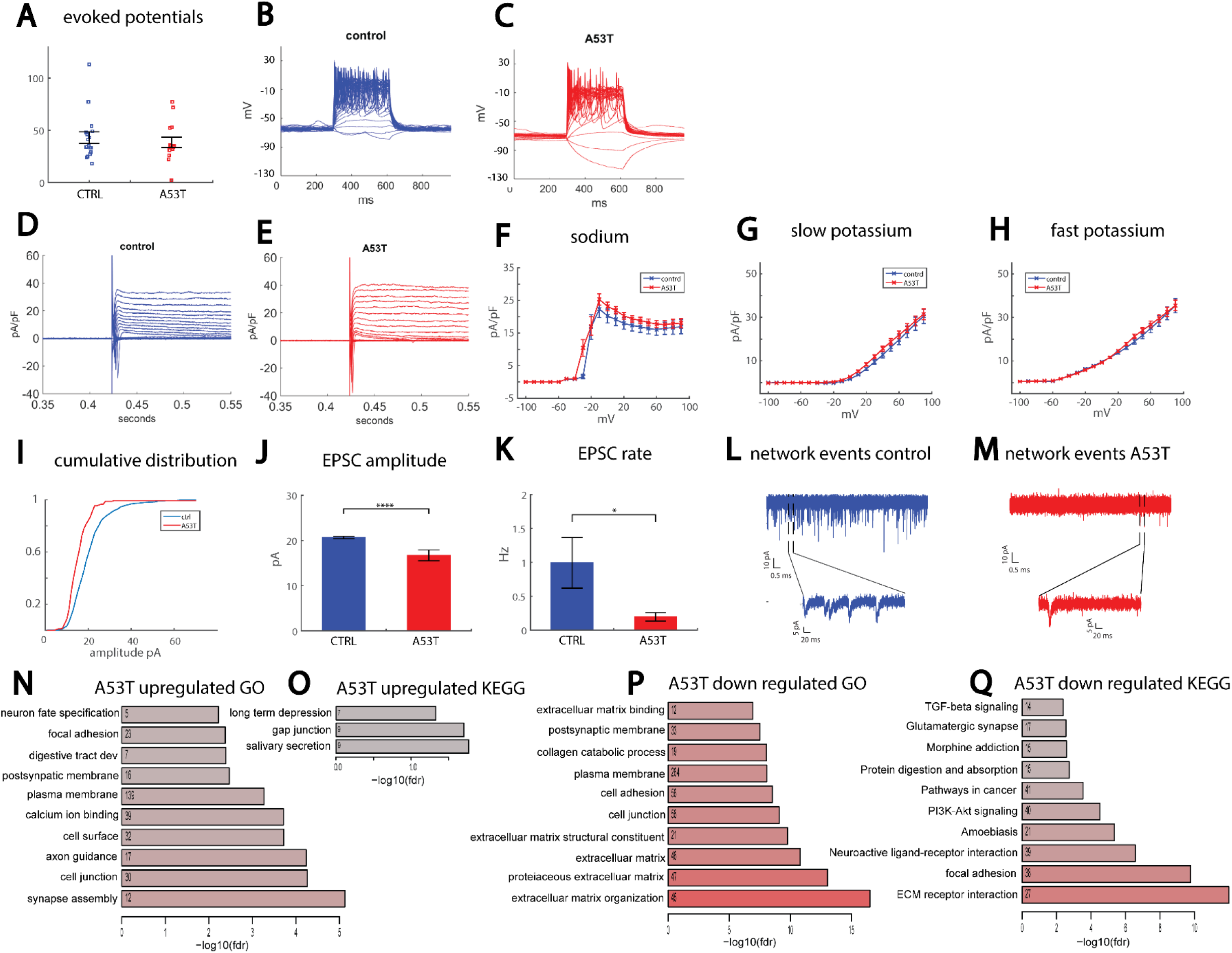
Reduced synaptic activity in neurons derived from an engineered line of iPSCs with an A53T mutation in the SNCA gene compared to isogenic controls. A. No significant change in the excitability of the A53T neurons compared to control neurons. B. Example recording of action potentials in current-clamp mode of a control neuron. C. Example recording of action potentials in current-clamp mode of an A53T neuron. Sodium currents of A53T neurons open at a lower depolarization potential than control neurons. D. Example recording of sodium and potassium currents in voltage-clamp mode in a control neuron. E. Example recording of sodium and potassium currents in voltage-clamp mode in an A53T neuron. F. No significant changes are observed in the sodium currents of the A53T neurons compared to controls. G. No significant changes are observed in the slow potassium currents in the A53T neurons. H. No significant changes are observed in the fast potassium currents in the A53T neurons. I. The cumulative distribution of the amplitudes of EPSCs is left-shifted in A53T neurons compared to control neurons, indicating lower amplitudes in the A53T neurons. J. The average amplitude of synaptic currents is significantly reduced in A53T neurons compared to control neurons. K. The average rate of synaptic currents is significantly reduced in A53T neurons compared to control neurons. L. Representative trace of the synaptic currents in a control neuron (the lower plot presents a zoom-in on the segment denoted in the black dashed lines in the upper graph). M. Representative trace of the synaptic currents in an A53T neuron (the lower plot presents a zoom-in on the segment denoted in the black dashed lines in the upper graph). N-Q. Bar plots showing top overrepresented terms or pathways among differential gene sets. N. Up-regulated GO terms in the A53T neurons compared to control neurons. O. Up-regulated KEGG pathways in DA A53T neurons compared to control neurons. P. Down-regulated GO terms in the A53T neurons compared to control neurons. Q. Down-regulated KEGG pathways in A53T neurons compared to control neurons. * p value<0.05, **** p values<0.0001. Bars represent the standard error in this figure.

The top 10 affected KEGG and MsigDB pathways and GO terms are presented in Figure 5N-Q, and the full list is given in Supplementary Figure 2-4. There were many dysregulated synapse-related pathways such as dopaminergic synapse, postsynaptic membrane, presynaptic membrane, chemical synaptic transmission, and more. Similar to the other PD lines presented in this study, there were many extracellular matrix-related affected pathways as well as pathways related to focal and cell adhesion, collagen processes, protein digestion and absorption, and PI3K-Akt signaling pathways.

### Protein aggregates and altered morphology in the A53T SNCA mutated iDopaNeurons compared to control neurons

We next delved deeper to look for additional morphological and cellular alterations that occurred in the engineered A53T iDopaNeurons neurons compared to controls. The neurons were stained for neuronal markers TUJ1 and MAP2, and both A53T and control neurons expressed these markers (Figure 6A-6D). Misfolded protein aggregates were observed in A53T neurons and rarely seen in control neurons (Figure. 6E-6J). The number of aggregates was quantified using the aggresomal kit and imaging in ultra-high resolution, 60.8 ± 7.1 % of the A53T neurons displayed protein aggregates (n=99), compared to 2.2±0.6 % of the control neurons (n=96, p<0.0001, Figure. 6Q). To assess the number of synapses, we co-stained for synapsin1 and the post-synaptic marker PSD95. Quantification of the number of puncta pairs Syn1/PSD95 density was significantly reduced in A53T cultures; 2.3±0.2 pairs in 10 μm were observed in n=15 neurites in control cultures, whereas only 1.5±0.16 (n=15 neurites) pairs in 10 μm were observed in A53T cultures (p=0.009, Figure 6K-P, Figure 6R). To assess how many of the misfolded protein aggregates were α-synuclein positive, we co-stained α-synuclein/aggresomes in the A53T neurons. Some of the aggregates contained α-synuclein protein (Figure 6S-U).

**Figure 6.**
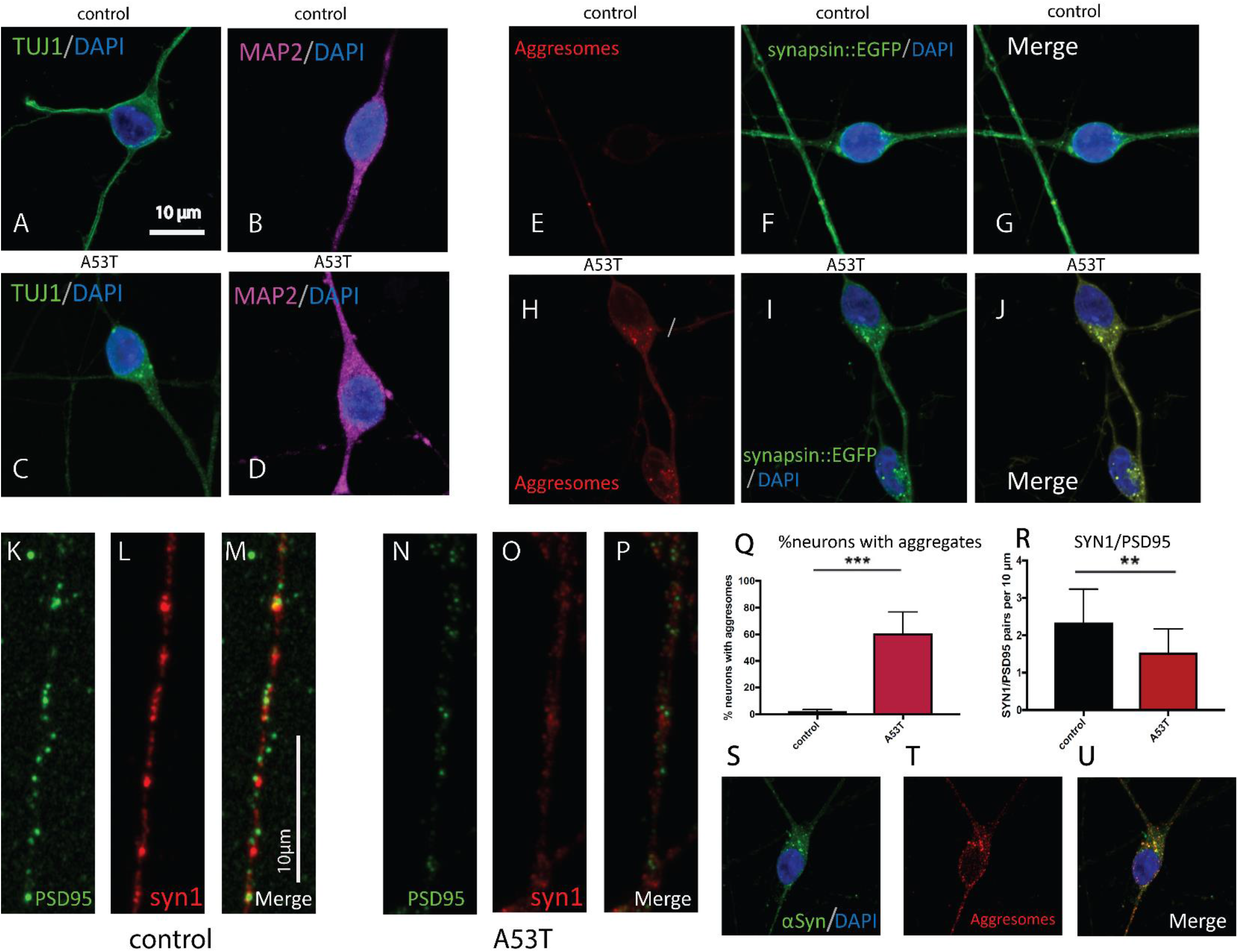
Protein aggregates in A53T dopaminergic neurons. A. Representative image showing expression of neuron-specific class III beta-tubulin/ 4’,6-diamidino-2-phenylindole (TUJ1/DAPI) in control dopaminergic neurons. Scale bar 10 μm. B. MAP2/DAPI expression in control dopaminergic neurons. C-D. Similar representative images of A53T dopaminergic neurons expressing TUJI/DAPI (C) and MAP2/DAPI (D). E-G. Control dopaminergic neurons exhibit almost no protein aggregates, as can be seen in the aggresomes and merged image with synapsin::EGFP/DAPI on the right. H-J. A53T dopaminergic neurons display protein aggregates, displayed in the aggresomes and synapsin::EGFP/DAPI staining. K-P. Immunostaining for postsynaptic density protein 95 (PSD95 in green) and synapsin1 (syn1) reveals a drastic decrease in the ratio of synapsin1 to PSD95, indicating fewer synapses. Q. Quantification of protein aggregates shows a drastic increase in the number of neurons harboring protein aggregates in the A53T mutated neurons compared to controls. R. Quantification of the ratio of syn1/PSD95 shows a drastic and significant decrease in A53T dopaminergic neurons compared to controls. S-U. α-Synuclein staining and aggresomes reveal co-localization of some of the protein aggregated with α-synuclein. ** p value<0.01, *** p values<0.001. Error bars represent standard deviations in this figure.

### Seeking commonly affected pathways in PD neurons

PD patients share similar symptoms despite completely different genes that cause the disease, or even when there is no defined genetic cause. Therefore, we sought to find common pathways between neurons derived from PD patients with different mutations. We started by pooling the RNA sequencing results from all our lines with PD-causing mutations - SNCA duplication, SNCA triplication, LRRK2, and Parkin - and we looked for differential expression compared to the 4 control lines. The 10 top terms enriched among up and down-regulated genes are presented in Figure 7A-D (GO and KEGG pathways). The full list as well as MsigDB dysregulated pathways are given in Supplementary Figure 2-4. A clear picture emerges of pathways that were strongly affected in neurons derived from all PD lines with mutations; some of them were not previously known to be associated with PD. Collagen-related pathways and the extracellular matrix were very strongly affected in the PD lines. Focal adhesion was another pathway that repeated throughout the monogenic PD lines, as well as the PI3K-Akt signaling pathway, pathways related to cancer and oxidoreductase activity, and protein digestion and absorption. Importantly, synapse-related pathways such as synapse, synaptic membrane, postsynapse, and more were commonly dysregulated in our monogenic PD lines. In the up-regulated pathways, we found cell adhesion molecules (CAM), which are molecules that interact with the extracellular matrix and may be a compensation mechanism for the reduced collagen and other extracellular matrix-related genes. We also looked for overrepresented GO terms for genes that were common in the monogenic PD lines and the sPD lines and these are presented in Figures 7E and 7F. The extracellular matrix is commonly implicated in both monogenic and sPD. Other dysregulated pathways were collagen pathways, focal and cell adhesion, PI3K-Akt signaling pathway, protein digestion and absorption, and a few pathways related to hypoxia. Pathways that were related to oxidative stress such as reactive oxygen species and oxidative stress were also dysregulated. Age-related pathways such as brain up, Alzheimer’s disease up, and cellular senescence were also commonly dysregulated in both monogenic and our sPD lines. Pathways that might be related to synapse function that were dysregulated included cell-cell junction, axon guidance, cell junction assembly, vasculature development, and regulation of neuron development projection.

**Figure 7.**
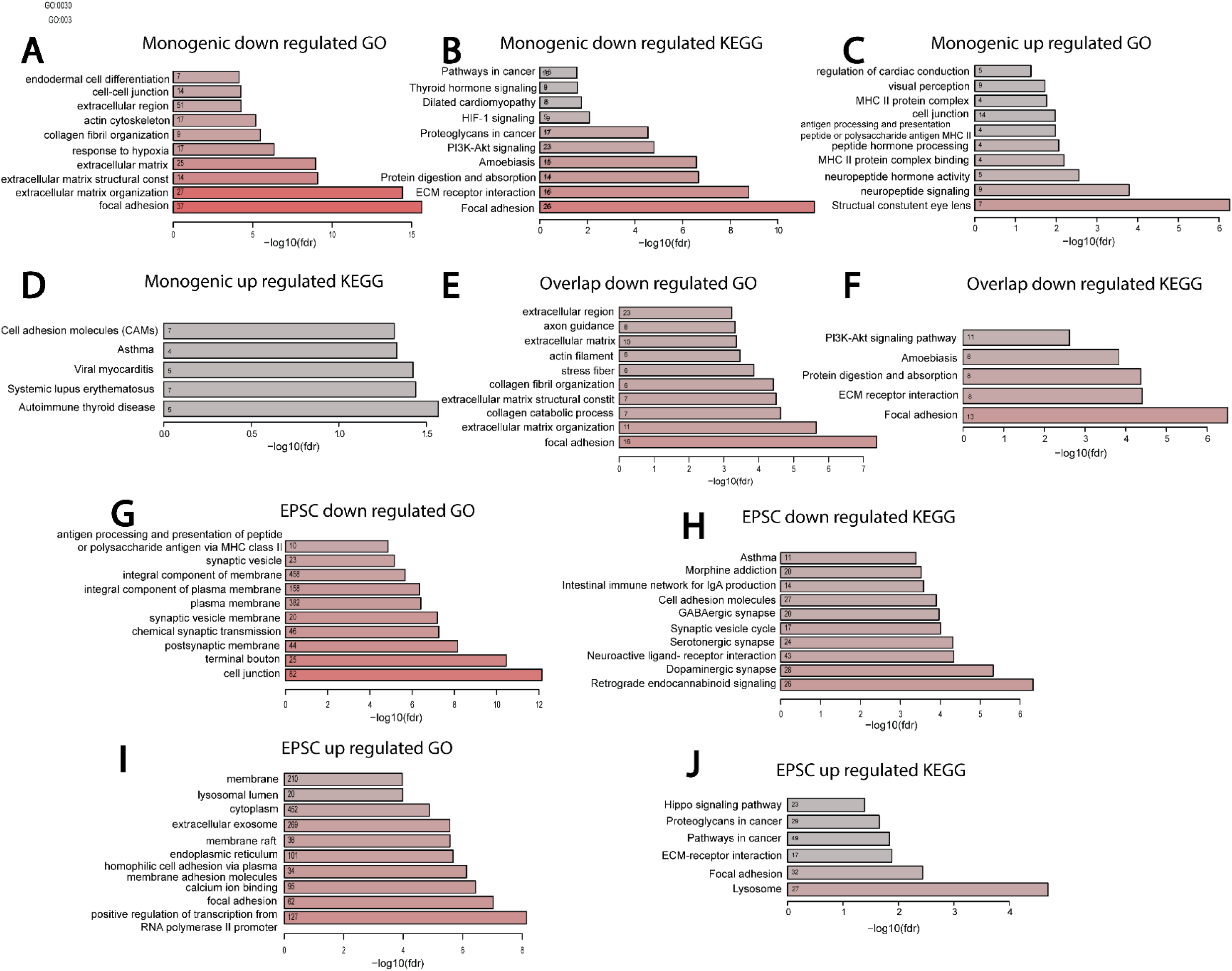
Pathways affected in monogenic (samples of all mutations pooled together) PD and pathways that are affected in neurons with a low rate of EPSCs. A. Top down-regulated GO terms in the monogenic PD compared to controls. B. Top down-regulated KEGG pathways in monogenic PD compared to controls. C. Top up-regulated GO terms in the monogenic PD compared to controls. D. Top up-regulated KEGG pathways in monogenic PD compared to controls. GO terms for up-regulated genes in the monogenic PD compared to controls. E. Commonly down-regulated GO terms in the monogenic neurons and the sPD neurons compared to the control neurons. F. Commonly down-regulated KEGG pathways in the monogenic neurons and the sPD neurons compared to the control neurons. G. Down-regulated GO terms in DA neurons that have a low rate of EPSCs. H. Down-regulated KEGG pathways in DA neurons that have a low rate of EPSCs. I. Up-regulated GO terms in DA neurons that have a low rate of EPSCs. J. Up-regulated KEGG pathways in DA neurons that have a low rate of EPSCs.

### Seeking pathways related directly to a reduction in synaptic transmission rate

The common electrophysiological phenotype that we observed in all PD lines was a drastic and significant reduction in the rate of synaptic current events. Therefore, we were interested to determine the affected pathways when the differential expression was taken relative to the EPSC rate of the investigated cell line (see Methods). The 10 top up-regulated pathways for neurons with reduced EPSC rates are presented in Figure 7G-J and the entire list of dysregulated GO terms and KEGG and MsigDB pathways is presented in Supplementary Figures 2-4. As expected, many synaptic pathways were dysregulated when comparing neurons that had a high-frequency rate of synaptic activity vs. neurons with a low rate of synaptic events. These included, for example, dopaminergic synapse, glutamatergic synapse, GABAergic synapse, serotonergic synapse, aminergic neurotransmitter loading into synaptic vesicles, filopodium, axon guidance, and many more. Other dysregulated pathways were similar to those that were dysregulated in PD, probably since PD lines exhibited a reduced synaptic event rate. These included many extracellular matrix pathways, focal adhesion, CAMs, melanosome, lysosome, cilium, mitochondrial pathways, endoplasmic reticulum-related pathways, and several cancer-related pathways.

## Discussion

PD affects the lives of nearly 1 million people in the US and is the most prevalent movement disorder. The hallmarks of PD are aggregates of the α-synuclein protein that are more specific to areas in the brain with a high density of DA neurons. In some PD cases, Lewy bodies appear in those DA-dense areas. These Lewy bodies are composed of protein aggregates whose main component is the α-synuclein protein. However, it is not clear if the Lewy bodies are causing the neuronal cell death that is observed in high-density DA neuron areas or are a side effect of other processes that occur and are the actual triggers for neurodegeneration. Recent studies suggest that degeneration starts as micro aggregates of the α-synuclein protein form, long before Lewy bodies appear. Strikingly, the PD-causing mutations are very different from one another, but most of the patients do not have a known genetic origin and are considered idiopathic or sporadic. Finding phenotypes that are common to PD early in the lifetime of neurons (and the patients) will help in deciphering disease mechanisms and finding effective treatment. Furthermore, the use of patient-derived neurons helps to mitigate the problem of the lack of animal models for sPD.

Here we report a neurophysiological phenotype that is common to DA neurons derived from PD patients for the following mutations: the A53T SNCA mutation, SNCA copy number variations (duplication as well as triplication), LRRK2, Parkin, and importantly, sPD patients. Using whole-cell patch-clamp, we found that neurons derived from PD patients all exhibited a significant reduction in the rate of synaptic activity. It is known that neurons that are plated sparsely have a low survival rate ^93, 94^; neurons need interactions with their neighboring neurons for survival. Therefore, the low connectivity that we observed in the PD neurons may exacerbate neuronal death caused by different mechanisms. Several labs have shown that reprogramming of adult cells into iPSCs erases aging signatures and epigenetic modifications ^76, 77^, and therefore the neurons in our cultures are considered to be young, even pre-natal neurons. Consequently, the observed phenotype of a reduction in synaptic activity in these very young neurons indicates an early process and a predisposition that is present in the patients’ DA neurons, probably long before patients exhibit any motor deficits. Similar findings were described in SNCA mutations in mice ^42, 43, 45, 47, 48^. In these mice, the earliest phenotype that was observed was a reduction in synaptic activity, followed by protein aggregates that, after weeks and sometimes months, evolved into the loss and death of DA neurons and to motor dysfunction. Overall, our results imply that neuronal cultures derived from human patients with the iPSC technique are a good model for studying PD progression, starting with this prodromal phenotype in the neurons. Moreover, this phenotype may have important implications in the early diagnosis and prediction of disease onset.

Even more remarkable, consistent with, and supportive of the electrophysiology were the shared dysregulated pathways that we revealed by analyzing gene expression using RNA sequencing. We observed dysregulation of pathways that were common in the different PD mutations as well as in the sPD patients. Among these dysregulated pathways and genes, we found the CAMs. Several CAM families have been shown to localize at the synapse ^95, 96^ and to influence the assembly and function of synapses in the CNS. The PI3K-Akt signaling pathways were also overrepresented among downregulated genes in both monogenic and sPD, and they have been shown to recruit PSD-95 to synapses ^97^. The FoxO pathway, another commonly overrepresented pathway among downregulated genes in our PD lines, has also been shown to play an important role in synaptic growth, synaptic vesicle exocytosis ^98^, and the promotion of synaptic plasticity ^99^. The latter two pathways have been shown to interact ^100^. Proteoglycans and collagen fibers also exhibited dysregulation in both our monogenic and sPD lines. These make up the extracellular matrix, which more and more evidence suggests is a part of the tetrapartite synapse. The extracellular matrix-related pathways were strongly dysregulated in almost all our PD lines. Overall, we saw dysregulation of many genes and pathways that affected synaptic formation. We hypothesize that this gene expression dosage dysregulation prevents neurons derived from PD patients from establishing effective synapses, measured in our electrophysiological experiments as a reduced rate of synaptic events. We hypothesize that PD neurons have a weakened ability to form operative synapses, which reduces the rate of synaptic events and triggers homeostatic mechanisms that take place; a vast dysregulation of genes associated with synaptic transmission-related pathways then occurs, as we observed when analyzing gene expression in our PD lines. It is interesting to note that, in those PD lines where there was a stronger decrease in the synaptic event’s rate, we also observed even more abundant synaptic-related pathways that were overrepresented among up-regulated genes. In the LRRK2 and the SNCA duplication, where the reduction in the synaptic rate was not as severe, we detected fewer synaptic pathways that were significantly enriched among up-regulated genes, further stressing the importance of electrophysiology for the detection of more subtle changes. Overall, our results strongly suggest that PD pathology starts at the synapse (as our in vitro neurons are young, and therefore reflect early brain events), in agreement with previous reports and hypotheses regarding the role of disrupted synapses in PD ^27, 101, 102^. Studies have shown that loss of synaptic terminals exceeds the loss of DA cell bodies ^74^ and that α-synuclein aggregates at the presynaptic terminals before forming Lewy bodies ^18, 28^; these results are supported by our findings of young and rejuvenated DA neurons that already exhibit synaptic deficits.

To summarize, our work demonstrates an early phenotype that is common to neurons from several PD mutations as well as sPD patients. It also demonstrates that there are common pathways that are affected in and common to PD patients (mutation-driven or sporadic). Importantly, these findings reveal that PD can be studied using iPSC-derived neuron technology, allowing us to trace the disease progression step by step. The affected pathways that we have identified through the analysis of gene expression should now be considered as important targets for further research.

## Acknowledgments

The authors would like to thank K.E. Diffenderfer for technical assistance and M.L. Gage for editorial comments. They would also like to acknowledge the Salk Institute Stem Cell Core, Waitt Biophotonics Core, and Next Generation Sequencing Core for technical support. Funding to the cores provided in part by NIH-NCI CCSG: P30 014195.

## Author contribution

SS differentiated neurons, did patch clamp experiments, performed data analysis, design of experiments, and MS writing, SL differentiated neurons prepared RNA, AM performed immunohistochemistry (IHC), imaging of aggregated proteins, MP differentiated neurons and performed IHC, IR differentiated neurons and performed IHC, MNS performed RNA sequencing analysis, FQ performed RNA sequencing analysis, SS performed RNA sequencing analysis, AM performed IHC, KM performed MEA experiments, TS performed confocal imaging and quantification, PO differentiated neurons, YS performed confocal imaging, AMD grew cells, LRM prepared lenti-virus, RN differentiated neurons, AA performed quantification of images, AR and TLW performed quantification of aggresome images, TN differentiated neurons, SBL performed RNA sequencing analysis, BW reprogrammed fibroblasts into iPSCs, BCF differentiated neurons, EJ helped with the design of experiments, CB performed path clamp experiments, AB diagnosed patients, JW diagnosed patients, MCM differentiated neurons and helped with design of experiments, FHG designed research and MS writing.

## Methods

### Ethics

The study was approved by the Salk institute with the following approvals: IRB 09-0003 and ESCRO 07-009.

### Human patients

Table 1 and 2 present the clinical features of the human patients who participated in this study. The first cohort, listed in Table 1, was diagnosed by Dr. Juergen Winkler, and the second cohort (Table 2) was diagnosed by Dr. Alexis Brice. A written consent was provided by all the participants in the study to take part of the study.

**Table 1.**
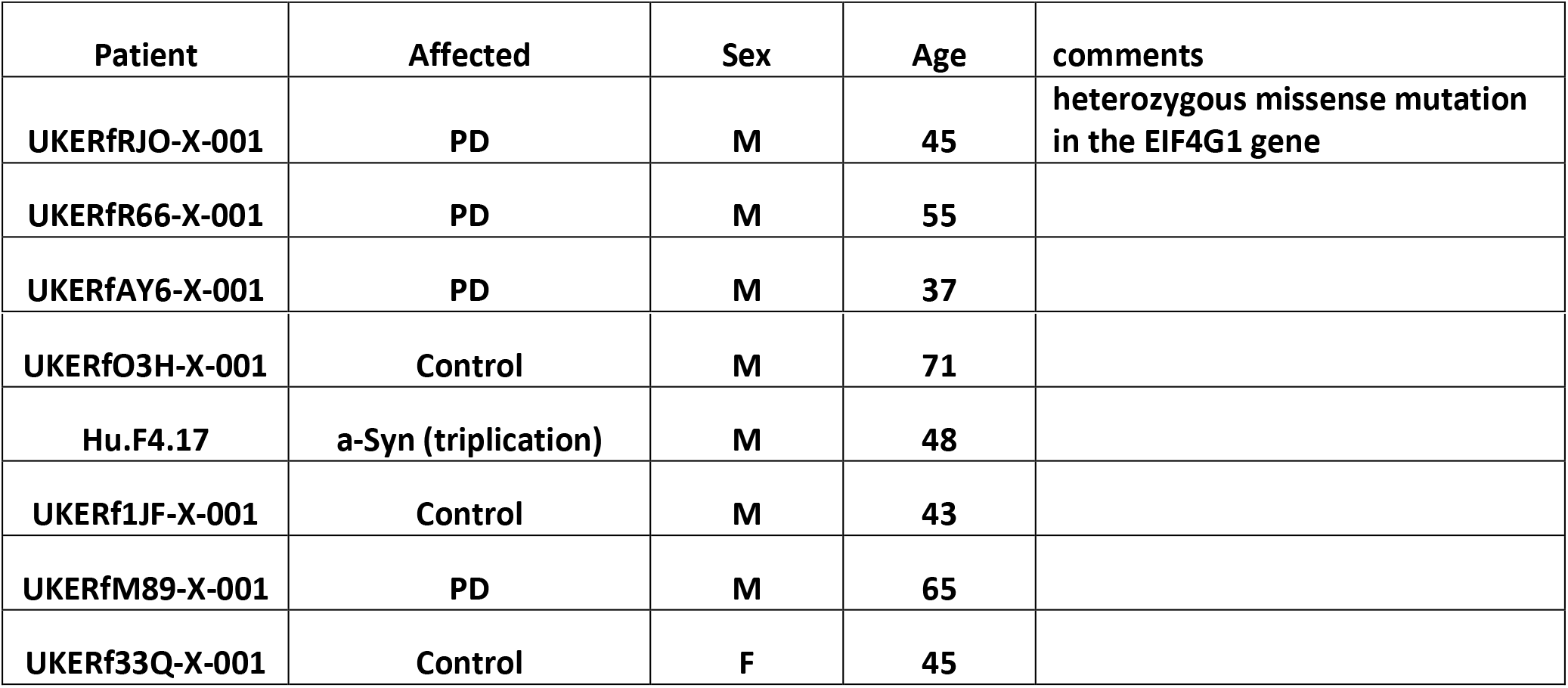
A description of the first cohort of the patients.

| Patient | Affected | Sex | Age | comments |
| --- | --- | --- | --- | --- |
| UKERfRJO-X-001 | PD | M | 45 | heterozygous missense mutation in the EIF4G1 gene |
| UKERfR66-X-001 | PD | M | 55 |  |
| UKERfAY6-X-001 | PD | M | 37 |  |
| **UKERfO3H-X-001** | **Control** | **M** | **71** |  |
| **Hu.F4.17** | **a-Syn (triplication)** | **M** | **48** |  |
| **UKERf1JF-X-001** | **Control** | **M** | **43** |  |
| **UKERfM89-X-001** | **PD** | **M** | **65** |  |
| **UKERf33Q-X-001** | **Control** | **F** | **45** |  |

**Table 2.**
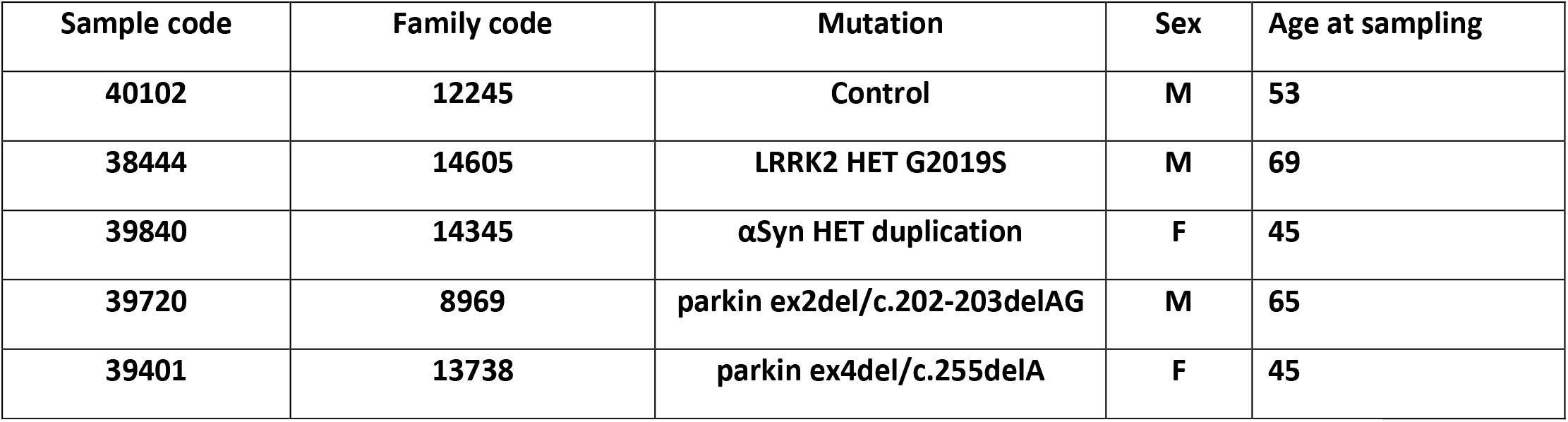
A description of the second cohort of the patients.

| **Sample code** | **Family code** | **Mutation** | **Sex** | **Age at sampling** |
| --- | --- | --- | --- | --- |
| **40102** | **12245** | **Control** | **M** | **53** |
| **38444** | **14605** | **LRRK2 HET G2019S** | **M** | **69** |
| **39840** | **14345** | **αSyn HET duplication** | **F** | **45** |
| **39720** | **8969** | **parkin ex2del/c.202-203delAG** | **M** | **65** |
| **39401** | **13738** | **parkin ex4del/c.255delA** | **F** | **45** |

### Dopaminergic (DA) neurons

DA neurons were generated from iPSCs based on a previously described protocol ^78^ with modifications ^79^. This protocol robustly produces around 70% DA neurons within the neuronal population in the culture. In brief, iPSCs were dissociated into a single cell suspension with TrypLE and replated on matrigel-coated plates at a density of 40,000 cells/cm^2^ in mTesR medium (Stem Cell Technologies). Cells were allowed to propagate with a daily change of medium for 2 days. Two days after the propagation of the cells, the differentiation process was started by switching into KSR medium (DMEM F-12 with Glutamax, 15% KO-SR, 1% NEAA, 1% Antibiotic-Antimycotic, 0.1 mM b-mercaptoethanol); this day was regarded as day 0. The medium was gradually changed to N2 medium (DMEM F-12 with Glutamax, 1% N2 supplement, 1% Antibiotic-Antimycotic) from day 5 to day 10 (day 5 and 6: 75% KSR : 25% N2; day 7 and 8: 50% KSR : 50% N2; day 9 and 10: 25% KSR : 75% N2). From Day 11 onward, the medium was switched to B27 medium (Neurobasal medium, 2% B27 supplement, 1% glutamax, 1% Antibiotic-Antimycotic, 10 ng/mL BDNF, 10 ng/mL GDNF, 1 ng/mL TGF3, 0.2 mM ascorbic acid and 0.1 mM cAMP). Cocktails of small molecules were added to the culture throughout the differentiation process (10 μM SB431542 on day 0-4; 100 nM LDN-193189 on day 0-12; 2 μM purmorphamine, 0.25 μM SAG, 100 ng/mL FGF8 on day 1-6; 3 μM CHIR99021 on day 3-12). Cells were replated onto Matrigel-coated coverslips on day 20 and further matured in B27 medium until day 30. On day 30, the base medium was switched to BrainPhys medium ^80^ until, on day 45-50, whole-cell patch-clamp recording was performed. All neurons that had a good quality patch-clamp seal were included in the analysis. The neurons in one of the experiments (results presented in Fig. 4) were grown and differentiated at the Bardy lab, using a published protocol ^80, 81^. This protocol produces a smaller proportion of tyrosine hydroxylase (TH)-positive neurons. Furthermore, the analysis was performed only on neurons that were classified as “type 5” neurons by this protocol.

### Plating neurons from a commercial control line and an affected line

We utilized commercial neurons from Fujifilm Cellular Dynamics International (CDI), which are a more homogeneous culture of neurons and produce more than 80% pure midbrain DA neurons The detailed nomenclatures/catalog numbers for the cells used in this paper are iCellͰDopaNeurons, 01279, catalog number: C1028; and MyCelliDopaNeurons, C1279, catalog number: C1113, genotype: SNCA (A53T alfa synuclein mutation). The cells were defrosted according to the protocol on the User’s Guide from CDI and counted. The medium used was Brainphys (Stem Cell Technologies), with the addition of supplements, according to the manufacturer’s instructions: iCellͰ Neural Supplement B (catalog number: M1029) and iCellͰ Nervous System Supplement (catalog number: M1031), N2 supplement, Laminin, and antibiotics (penicillin/streptomycin). For patch clamp electrophysiology experiments, we used Poly-L-ornithine- and Laminin-coated, 24-well plates with coverslips in each well, and we plated 4-6×10^5^ cells per well. Recordings started after a week in culture. For the RNA sequencing experiments, we plated 1×10^6^ cells per well on 6-well plates. For immunostaining, we plated about 1×10^5^ cells per well of 8 chambers ibid_Ͱ_ slides.

### iCell DopaNeuron cultures on multi-electrode arrays (MEAs)

Control iCell DopaNeurons (Fujifilm Cellular Dynamics Inc.) and MyCell *SNCA* A53T DopaNeurons were thawed and dotted (~80k cells) onto a 48-well MEA plate (Axion Biosystems) using the iCell DopaNeuron MEA application protocol. Cultures were treated with BrainPhys medium (Stem Cell Technologies), with 50% medium changes performed every 2 to 3 days for 20 days. Five-minute recordings were made on DIV20 post-plating (N=24). Raw voltage recordings were processed via a Butterworth (200-4kHz) filter; action potentials were then detected by a 5.5 standard deviation detection threshold, and network-level bursting behaviors were analyzed off-line using the Axion’s NeuralMetric toolbox via the ‘Envelope’ algorithm (threshold factor 3, minimum inter-burst-interval 10 seconds, burst inclusion percentage 75, minimum number of electrodes percentage 25) and synchrony index (20 milliseconds) ^48^. The activity and network-level bursting behaviors assessed included mean firing rate, network bursting rate (bursts per minute: BPM), intensity (Hz), and duration (seconds). A two-tailed student t-test was used to assess statistical differences on each measure independently between the 2 groups of neurons.

### Electrophysiology – Whole-cell patch-clamp

Neurons on glass coverslips were transferred to a recording chamber in a standard recording medium containing (in mM) 10 HEPES, 4 KCl, 2 CaCl_2_, 1 MgCl_2_, 139 NaCl, 10 D-glucose (310 mOsm, pH 7.4). Whole-cell patch-clamp recordings were performed on days 9-12 days after plating for A53T vs. iDOPA CDI lines and for 40 days after plating for the other DA protocols. Patch electrodes were filled with internal solutions containing (in mM) 130 K-gluconate, 6 KCl, 4 NaCl, 10 Na-HEPES, 0.2 K-EGTA, 0.3 GTP, 2 Mg-ATP, 0.2 cAMP, 10 D-glucose, 0.15% biocytin and 0.06% rhodamine. The pH and osmolarity of the internal solution were brought close to physiological conditions (pH 7.3, 290–300 mOsmol). Signals were amplified with a Multiclamp700B amplifier and recorded with Clampex 10.2 software (Axon Instruments). Data were acquired at a sampling rate of 20 kHz and analyzed using Clampfit-10 and the software package Matlab (2018b, The MathWorks Inc., Natick, MA, 2000). All measurements were conducted at room temperature. For post-synaptic current measurements, 40 μM of bicuculline was applied and cells were held at −60 mV when currents were recorded (−70 mV after correction of junction liquid potential).

### Analysis electrophysiology

#### Total evoked action potentials

Cells were typically held in current-clamp mode near −60 mV with a steady holding current, and current injections were given starting 12 pA below the steady holding current in 3-pA steps 400 ms in duration. A total of 35 depolarization steps were injected in current-clamp mode. Neurons that needed more than 50 pA to be held at −60 mV were discarded from the analysis. The total number of action potentials was counted in the first 17 depolarization steps.

#### Action potential shape analysis

The first evoked action potential was used for spike shape analysis (with the lowest injected current needed for eliciting an action potential). Spike threshold was the membrane potential at which the slope of the depolarizing membrane potential increased drastically, resulting in an action potential (the first maximum in the second derivative of the voltage vs. time). The fast (5 ms) AHP amplitude was calculated as the difference between the threshold of the action potential and the value of the membrane potential 5 ms after the potential returned to cross the threshold value at the end of the action potential. The spike amplitude was calculated as the difference between the maximum membrane potential during a spike and the threshold. Action potential width was calculated as the time it took for the membrane potential to reach half the spike amplitude in the rising part of the spike to the descending part of the spike (Full Width at Half Maximum).

#### Input conductance

The input conductance was calculated around the resting membrane potential by measuring the current with the cell held in voltage-clamp mode first at −70 mV and then at −50 mV. The difference in currents divided by the difference in membrane potential (of 20 mV) is the calculated input conductance.

#### Sodium and potassium currents

The sodium and potassium currents were acquired in voltage-clamp mode. Cells were held at −60 mV, and voltage steps of 400 ms were made in the range of −90 mV to 80 mV. Currents were normalized by the cell capacitance.

#### Fast and slow potassium currents

We measured the fast potassium current by the maximum current immediately following a depolarization step, typically within a time window of a few milliseconds. The slow potassium currents were obtained at the end of the 400-ms depolarization step.

#### Capacitance

The capacitance was measured by the membrane test of the Clampex SW.

#### Synaptic activity

Excitatory synaptic activity was measured in voltage-clamp mode with 40 μM bicuculline applied in the recording medium. The neurons were held at −60 mV and currents were measured in the patched neuron. We measured both the amplitude of these currents and their rates. A neuron was defined as having synaptic activity if it had more than 20 events in a 60-second recording and as not having synaptic activity if it had fewer than 20 events in 60 seconds.

### RNA-Sequencing analysis

Sequenced reads were quality-tested using FASTQC ^82^ v0.11.5 and aligned to the hg19 ^83^ human genome using the STAR aligner ^84^ version 2.5.3a. Mapping was carried out using default parameters, filtering non-canonical introns and allowing up to 10 mismatches per read, and only keeping uniquely mapped reads. The genome index was constructed using the gene annotation supplied with the hg19 Illumina iGenomes collection ^85^ and sjdbOverhang value of 100. Raw or TPM (Transcripts per million) gene expression was quantified across all gene exons with HOMER ^86^ using the top-expressed isoform as a proxy for gene expression, and differential gene expression was carried out on the raw counts using the edgeR ^87^ package version 3.28.1. For each disease type, differentially expressed genes were defined as having a false discovery rate (FDR) <0.05 when comparing 2 experimental conditions. We also separated monogenic subjects from sPD and compared each group separately to controls using the exact test function in edgeR. We then combined all subjects (monogenic, sPD, and control) and treated the EPSC rate as a continuous variable; we conducted differential expression analysis using the GLM model in edgeR. In both cases, differentially expressed genes were defined as having an FDR < 0.05. A GO enrichment test and KEGG pathway analysis were performed using the program DAVID Bioinformatics Resources 6.8 ^88^. Overrepresentation of GO terms and KEGG pathway was determined by FDR < 0.05. MsigDB ^89^ overrepresentation analysis was carried out using HOMER findGO.pl using the corrected Benjamini & Yakutieli method for multiple testing correction ^90^.

### Immunocytochemistry and cell imaging

Neuronal cultures were fixed with 4% paraformaldehyde for 15 minutes at room temperature and then treated with PBS containing 0.1% Triton X-100. After a 15-minute PBS wash, cells were blocked with 5% BSA in PBS for 1 hour, then incubated with the primary antibody in PBS at 4^0^C overnight and the next day after a few PBS washes with secondary antibodies for 1 hour at ambient temperature. This process was followed by a 10-minute incubation with DAPI and a final set of PBS washes. The coverslips were mounted on glass slides using PVA-DABCO. Primary antibodies used were rabbit anti-Tuj1, (1:500, Covance), chicken anti-MAP2 (1:400, Abcam), rabbit anti-αSynuclein (1:500, Invitrogen), rabbit anti-TH (1:500, Pel-Freez), mouse anti-PSD95 (1:500, Life Tech), rabbit anti-Synapsin I (1:500, Calbiochem). Corresponding Alexa Fluor™ secondary antibodies were then used (1:1000). For detecting protein aggregates, the PROTEOSTAT® Aggresome Detection kit was used according to the manufacturer’s instructions.

Confocal z-stacks were acquired with a Zeiss LSM 880 Airy scan microscope (Carl Zeiss, Microimaging Inc.) using 405 nm Diode laser, 488 nm Argon, and 543 nm HeNe lasers with a Plan NeoFluar 40X/1.3 oil DIC or a Plan-Apochromat 63X/1.4 oil DIC objective.

### Imaging of cell morphology

Neurons stained for TH were imaged and analyzed with Neurolucida SW (MBF Bioscience), where neurites and soma were manually traced.

### Statistical analysis

Unless otherwise stated, p values were calculated using a two-sample t-test (two-tailed).

## Data Availability

The datasets generated and analysed during the current study are available in the figshare repository in the following link: https://figshare.com/articles/dataset/ephys_data_7z/14635452.

## References

1. Parkinson J. An essay on the shaking palsy. 1817. The Journal of neuropsychiatry and clinical neurosciences 2002; 14(2):223–236; discussion 222.

2. Tysnes OB, Storstein A. Epidemiology of Parkinson’s disease. Journal of neural transmission 2017; 124(8):901–905.

3. von Campenhausen S, Bornschein B, Wick R, Botzel K, Sampaio C, Poewe W et al. Prevalence and incidence of Parkinson’s disease in Europe. European neuropsychopharmacology : the journal of the European College of Neuropsychopharmacology 2005; 15(4):473–490.

4. Bagheri H, Damase-Michel C, Lapeyre-Mestre M, Cismondo S, O’Connell D, Senard JM et al. A study of salivary secretion in Parkinson’s disease. Clinical neuropharmacology 1999; 22(4):213–215.

5. Huang P, Yang XD, Chen SD, Xiao Q. The association between Parkinson’s disease and melanoma: a systematic review and meta-analysis. Translational neurodegeneration 2015; 4:21.

6. Braak H, Del Tredici K, Rub U, de Vos RA, Jansen Steur EN, Braak E. Staging of brain pathology related to sporadic Parkinson’s disease. Neurobiology of aging 2003; 24(2):197–211.

7. Burke RE, Dauer WT, Vonsattel JP. A critical evaluation of the Braak staging scheme for Parkinson’s disease. Annals of neurology 2008; 64(5):485–491.

8. Jellinger KA. Critical evaluation of the Braak staging scheme for Parkinson’s disease. Annals of neurology 2010; 67(4):550.

9. Tran J, Anastacio H, Bardy C. Genetic predispositions of Parkinson’s disease revealed in patient-derived brain cells. NPJ Parkinsons Dis 2020; 6:8.

10. Li JQ, Tan L, Yu JT. The role of the LRRK2 gene in Parkinsonism. Mol Neurodegener 2014; 9:47.

11. Nuytemans K, Theuns J, Cruts M, Van Broeckhoven C. Genetic etiology of Parkinson disease associated with mutations in the SNCA, PARK2, PINK1, PARK7, and LRRK2 genes: a mutation update. Human mutation 2010; 31(7):763–780.

12. Cummings JL. The dementias of Parkinson’s disease: prevalence, characteristics, neurobiology, and comparison with dementia of the Alzheimer type. Eur Neurol 1988; 28 Suppl 1:15–23.

13. Chartier-Harlin MC, Kachergus J, Roumier C, Mouroux V, Douay X, Lincoln S et al. Alpha-synuclein locus duplication as a cause of familial Parkinson’s disease. Lancet 2004; 364(9440):1167–1169.

14. Golbe LI, Di Iorio G, Bonavita V, Miller DC, Duvoisin RC. A large kindred with autosomal dominant Parkinson’s disease. Annals of neurology 1990; 27(3):276–282.

15. Singleton AB, Farrer M, Johnson J, Singleton A, Hague S, Kachergus J et al. alpha-Synuclein locus triplication causes Parkinson’s disease. Science 2003; 302(5646):841.

16. Domingo A, Klein C. Genetics of Parkinson disease. Handb Clin Neurol 2018; 147:211–227.

17. Kramer ML, Schulz-Schaeffer WJ. Presynaptic alpha-synuclein aggregates, not Lewy bodies, cause neurodegeneration in dementia with Lewy bodies. The Journal of neuroscience : the official journal of the Society for Neuroscience 2007; 27(6):1405–1410.

18. Schulz-Schaeffer WJ. Is Cell Death Primary or Secondary in the Pathophysiology of Idiopathic Parkinson’s Disease? Biomolecules 2015; 5(3):1467–1479.

19. Tanaka M, Kim YM, Lee G, Junn E, Iwatsubo T, Mouradian MM. Aggresomes formed by alpha-synuclein and synphilin-1 are cytoprotective. The Journal of biological chemistry 2004; 279(6):4625–4631.

20. Luk KC, Kehm V, Carroll J, Zhang B, O’Brien P, Trojanowski JQ et al. Pathological alpha-synuclein transmission initiates Parkinson-like neurodegeneration in nontransgenic mice. Science 2012; 338(6109):949–953.

21. Nuber S, Rajsombath M, Minakaki G, Winkler J, Muller CP, Ericsson M et al. Abrogating Native alpha-Synuclein Tetramers in Mice Causes a L-DOPA-Responsive Motor Syndrome Closely Resembling Parkinson’s Disease. Neuron 2018; 100(1):75–90 e75.

22. Burre J. The Synaptic Function of alpha-Synuclein. J Parkinsons Dis 2015; 5(4):699–713.

23. George JM. The synucleins. Genome biology 2002; 3(1):REVIEWS3002.

24. Withers GS, George JM, Banker GA, Clayton DF. Delayed localization of synelfin (synuclein, NACP) to presynaptic terminals in cultured rat hippocampal neurons. Brain research Developmental brain research 1997; 99(1):87–94.

25. Stefanis L. alpha-Synuclein in Parkinson’s disease. Cold Spring Harbor perspectives in medicine 2012; 2(2):a009399.

26. Bridi JC, Hirth F. Mechanisms of alpha-Synuclein Induced Synaptopathy in Parkinson’s Disease. Front Neurosci 2018; 12:80.

27. Soukup SF, Vanhauwaert R, Verstreken P. Parkinson’s disease: convergence on synaptic homeostasis. EMBO J 2018; 37(18).

28. Spinelli KJ, Taylor JK, Osterberg VR, Churchill MJ, Pollock E, Moore C et al. Presynaptic alpha-synuclein aggregation in a mouse model of Parkinson’s disease. The Journal of neuroscience : the official journal of the Society for Neuroscience 2014; 34(6):2037–2050.

29. Fuchs J, Nilsson C, Kachergus J, Munz M, Larsson EM, Schule B et al. Phenotypic variation in a large Swedish pedigree due to SNCA duplication and triplication. Neurology 2007; 68(12):916–922.

30. Ki CS, Stavrou EF, Davanos N, Lee WY, Chung EJ, Kim JY et al. The Ala53Thr mutation in the alpha-synuclein gene in a Korean family with Parkinson disease. Clinical genetics 2007; 71(5):471–473.

31. Konno T, Ross OA, Puschmann A, Dickson DW, Wszolek ZK. Autosomal dominant Parkinson’s disease caused by SNCA duplications. Parkinsonism & related disorders 2016; 22 Suppl 1:S1–6.

32. Kruger R, Kuhn W, Muller T, Woitalla D, Graeber M, Kosel S et al. Ala30Pro mutation in the gene encoding alpha-synuclein in Parkinson’s disease. Nature genetics 1998; 18(2):106–108.

33. Polymeropoulos MH, Lavedan C, Leroy E, Ide SE, Dehejia A, Dutra A et al. Mutation in the alpha-synuclein gene identified in families with Parkinson’s disease. Science 1997; 276(5321):2045–2047.

34. Puschmann A, Ross OA, Vilarino-Guell C, Lincoln SJ, Kachergus JM, Cobb SA et al. A Swedish family with de novo alpha-synuclein A53T mutation: evidence for early cortical dysfunction. Parkinsonism & related disorders 2009; 15(9):627–632.

35. Zarranz JJ, Alegre J, Gomez-Esteban JC, Lezcano E, Ros R, Ampuero I et al. The new mutation, E46K, of alpha-synuclein causes Parkinson and Lewy body dementia. Annals of neurology 2004; 55(2):164–173.

36. Murphy DD, Rueter SM, Trojanowski JQ, Lee VM. Synucleins are developmentally expressed, and alpha-synuclein regulates the size of the presynaptic vesicular pool in primary hippocampal neurons. The Journal of neuroscience : the official journal of the Society for Neuroscience 2000; 20(9):3214–3220.

37. Scott D, Roy S. alpha-Synuclein inhibits intersynaptic vesicle mobility and maintains recycling-pool homeostasis. The Journal of neuroscience : the official journal of the Society for Neuroscience 2012; 32(30):10129–10135.

38. Liu S, Ninan I, Antonova I, Battaglia F, Trinchese F, Narasanna A et al. alpha-Synuclein produces a long-lasting increase in neurotransmitter release. EMBO J 2004; 23(22):4506–4516.

39. Lotharius J, Brundin P. Impaired dopamine storage resulting from alpha-synuclein mutations may contribute to the pathogenesis of Parkinson’s disease. Human molecular genetics 2002; 11(20):2395–2407.

40. Yavich L, Tanila H, Vepsalainen S, Jakala P. Role of alpha-synuclein in presynaptic dopamine recruitment. The Journal of neuroscience : the official journal of the Society for Neuroscience 2004; 24(49):11165–11170.

41. Eslamboli A, Romero-Ramos M, Burger C, Bjorklund T, Muzyczka N, Mandel RJ et al. Long-term consequences of human alpha-synuclein overexpression in the primate ventral midbrain. Brain : a journal of neurology 2007; 130(Pt 3):799–815.

42. Ip CW, Klaus LC, Karikari AA, Visanji NP, Brotchie JM, Lang AE et al. AAV1/2-induced overexpression of A53T-alpha-synuclein in the substantia nigra results in degeneration of the nigrostriatal system with Lewy-like pathology and motor impairment: a new mouse model for Parkinson’s disease. Acta neuropathologica communications 2017; 5(1):11.

43. Kirik D, Rosenblad C, Burger C, Lundberg C, Johansen TE, Muzyczka N et al. Parkinson-like neurodegeneration induced by targeted overexpression of alpha-synuclein in the nigrostriatal system. The Journal of neuroscience : the official journal of the Society for Neuroscience 2002; 22(7):2780–2791.

44. Kouroupi G, Taoufik E, Vlachos IS, Tsioras K, Antoniou N, Papastefanaki F et al. Defective synaptic connectivity and axonal neuropathology in a human iPSC-based model of familial Parkinson’s disease. Proceedings of the National Academy of Sciences of the United States of America 2017; 114(18):E3679–E3688.

45. Lundblad M, Decressac M, Mattsson B, Bjorklund A. Impaired neurotransmission caused by overexpression of alpha-synuclein in nigral dopamine neurons. Proceedings of the National Academy of Sciences of the United States of America 2012; 109(9):3213–3219.

46. Nemani VM, Lu W, Berge V, Nakamura K, Onoa B, Lee MK et al. Increased expression of alpha-synuclein reduces neurotransmitter release by inhibiting synaptic vesicle reclustering after endocytosis. Neuron 2010; 65(1):66–79.

47. Paumier KL, Sukoff Rizzo SJ, Berger Z, Chen Y, Gonzales C, Kaftan E et al. Behavioral characterization of A53T mice reveals early and late stage deficits related to Parkinson’s disease. PloS one 2013; 8(8):e70274.

48. Wu N, Joshi PR, Cepeda C, Masliah E, Levine MS. Alpha-synuclein overexpression in mice alters synaptic communication in the corticostriatal pathway. Journal of neuroscience research 2010; 88(8):1764–1776.

49. Fishbein I, Segal M. Miniature synaptic currents become neurotoxic to chronically silenced neurons. Cerebral cortex 2007; 17(6):1292–1306.

50. Bardien S, Lesage S, Brice A, Carr J. Genetic characteristics of leucine-rich repeat kinase 2 (LRRK2) associated Parkinson’s disease. Parkinsonism & related disorders 2011; 17(7):501–508.

51. Rideout HJ, Stefanis L. The neurobiology of LRRK2 and its role in the pathogenesis of Parkinson’s disease. Neurochem Res 2014; 39(3):576–592.

52. Venderova K, Kabbach G, Abdel-Messih E, Zhang Y, Parks RJ, Imai Y et al. Leucine-Rich Repeat Kinase 2 interacts with Parkin, DJ-1 and PINK-1 in a Drosophila melanogaster model of Parkinson’s disease. Human molecular genetics 2009; 18(22):4390–4404.

53. Liu Z, Wang X, Yu Y, Li X, Wang T, Jiang H et al. A Drosophila model for LRRK2-linked parkinsonism. Proceedings of the National Academy of Sciences of the United States of America 2008; 105(7):2693–2698.

54. Saha S, Guillily MD, Ferree A, Lanceta J, Chan D, Ghosh J et al. LRRK2 modulates vulnerability to mitochondrial dysfunction in Caenorhabditis elegans. The Journal of neuroscience : the official journal of the Society for Neuroscience 2009; 29(29):9210–9218.

55. Li Y, Liu W, Oo TF, Wang L, Tang Y, Jackson-Lewis V et al. Mutant LRRK2(R1441G) BAC transgenic mice recapitulate cardinal features of Parkinson’s disease. Nat Neurosci 2009; 12(7):826–828.

56. Tong Y, Pisani A, Martella G, Karouani M, Yamaguchi H, Pothos EN et al. R1441C mutation in LRRK2 impairs dopaminergic neurotransmission in mice. Proceedings of the National Academy of Sciences of the United States of America 2009; 106(34):14622–14627.

57. Cirnaru MD, Marte A, Belluzzi E, Russo I, Gabrielli M, Longo F et al. LRRK2 kinase activity regulates synaptic vesicle trafficking and neurotransmitter release through modulation of LRRK2 macro-molecular complex. Front Mol Neurosci 2014; 7:49.

58. Inoshita T, Arano T, Hosaka Y, Meng H, Umezaki Y, Kosugi S et al. Vps35 in cooperation with LRRK2 regulates synaptic vesicle endocytosis through the endosomal pathway in Drosophila. Human molecular genetics 2017; 26(15):2933–2948.

59. Shin N, Jeong H, Kwon J, Heo HY, Kwon JJ, Yun HJ et al. LRRK2 regulates synaptic vesicle endocytosis. Exp Cell Res 2008; 314(10):2055–2065.

60. Piccoli G, Condliffe SB, Bauer M, Giesert F, Boldt K, De Astis S et al. LRRK2 controls synaptic vesicle storage and mobilization within the recycling pool. The Journal of neuroscience : the official journal of the Society for Neuroscience 2011; 31(6):2225–2237.

61. Sassone J, Serratto G, Valtorta F, Silani V, Passafaro M, Ciammola A. The synaptic function of parkin. Brain : a journal of neurology 2017; 140(9):2265–2272.

62. Huttner WB, Schiebler W, Greengard P, De Camilli P. Synapsin I (protein I), a nerve terminal-specific phosphoprotein. III. Its association with synaptic vesicles studied in a highly purified synaptic vesicle preparation. J Cell Biol 1983; 96(5):1374–1388.

63. Kubo SI, Kitami T, Noda S, Shimura H, Uchiyama Y, Asakawa S et al. Parkin is associated with cellular vesicles. J Neurochem 2001; 78(1):42–54.

64. Fallon L, Moreau F, Croft BG, Labib N, Gu WJ, Fon EA. Parkin and CASK/LIN-2 associate via a PDZ-mediated interaction and are co-localized in lipid rafts and postsynaptic densities in brain. The Journal of biological chemistry 2002; 277(1):486–491.

65. Lucking CB, Durr A, Bonifati V, Vaughan J, De Michele G, Gasser T et al. Association between early-onset Parkinson’s disease and mutations in the parkin gene. N Engl J Med 2000; 342(21):1560–1567.

66. Pankratz N, Kissell DK, Pauciulo MW, Halter CA, Rudolph A, Pfeiffer RF et al. Parkin dosage mutations have greater pathogenicity in familial PD than simple sequence mutations. Neurology 2009; 73(4):279–286.

67. Pankratz N, Wilk JB, Latourelle JC, DeStefano AL, Halter C, Pugh EW et al. Genomewide association study for susceptibility genes contributing to familial Parkinson disease. Human genetics 2009; 124(6):593–605.

68. Poulopoulos M, Levy OA, Alcalay RN. The neuropathology of genetic Parkinson’s disease. Movement disorders : official journal of the Movement Disorder Society 2012; 27(7):831–842.

69. Helton TD, Otsuka T, Lee MC, Mu Y, Ehlers MD. Pruning and loss of excitatory synapses by the parkin ubiquitin ligase. Proceedings of the National Academy of Sciences of the United States of America 2008; 105(49):19492–19497.

70. Trempe JF, Chen CX, Grenier K, Camacho EM, Kozlov G, McPherson PS et al. SH3 domains from a subset of BAR proteins define a Ubl-binding domain and implicate parkin in synaptic ubiquitination. Molecular cell 2009; 36(6):1034–1047.

71. Coetzee SG, Pierce S, Brundin P, Brundin L, Hazelett DJ, Coetzee GA. Enrichment of risk SNPs in regulatory regions implicate diverse tissues in Parkinson’s disease etiology. Scientific reports 2016; 6:30509.

72. Zhu M, Cortese GP, Waites CL. Parkinson’s disease-linked Parkin mutations impair glutamatergic signaling in hippocampal neurons. BMC Biol 2018; 16(1):100.

73. Burke RE, O’Malley K. Axon degeneration in Parkinson’s disease. Experimental neurology 2013; 246:72–83.

74. Cheng HC, Ulane CM, Burke RE. Clinical progression in Parkinson disease and the neurobiology of axons. Annals of neurology 2010; 67(6):715–725.

75. Hornykiewicz O. Biochemical aspects of Parkinson’s disease. Neurology 1998; 51(2 Suppl 2):S2–9.

76. Mertens J, Paquola ACM, Ku M, Hatch E, Bohnke L, Ladjevardi S et al. Directly Reprogrammed Human Neurons Retain Aging-Associated Transcriptomic Signatures and Reveal Age-Related Nucleocytoplasmic Defects. Cell stem cell 2015; 17(6):705–718.

77. Rando TA, Chang HY. Aging, rejuvenation, and epigenetic reprogramming: resetting the aging clock. Cell 2012; 148(1-2):46–57.

78. Kriks S, Shim JW, Piao J, Ganat YM, Wakeman DR, Xie Z et al. Dopamine neurons derived from human ES cells efficiently engraft in animal models of Parkinson’s disease. Nature 2011; 480(7378):547–551.

79. Zhang P, Xia N, Reijo Pera RA. Directed dopaminergic neuron differentiation from human pluripotent stem cells. Journal of visualized experiments : JoVE 2014; (91):51737.

80. Bardy C, van den Hurk M, Eames T, Marchand C, Hernandez RV, Kellogg M et al. Neuronal medium that supports basic synaptic functions and activity of human neurons in vitro. Proceedings of the National Academy of Sciences of the United States of America 2015; 112(20):E2725–2734.

81. Bardy C, van den Hurk M, Kakaradov B, Erwin JA, Jaeger BN, Hernandez RV et al. Predicting the functional states of human iPSC-derived neurons with single-cell RNA-seq and electrophysiology. Mol Psychiatry 2016; 21(11):1573–1588.

82. Andrews S. A quality control tool for high throughput sequence data. Babraham Bioinformatics 2010.

83. Lander ES, Linton LM, Birren B, Nusbaum C, Zody MC, Baldwin J et al. Initial sequencing and analysis of the human genome. Nature 2001; 409(6822):860–921.

84. Dobin A, Davis CA, Schlesinger F, Drenkow J, Zaleski C, Jha S et al. STAR: ultrafast universal RNA-seq aligner. Bioinformatics 2013; 29(1):15–21.

85. Illumina. iGenomes online 2015.

86. Heinz S, Benner C, Spann N, Bertolino E, Lin YC, Laslo P et al. Simple combinations of lineage-determining transcription factors prime cis-regulatory elements required for macrophage and B cell identities. Molecular cell 2010; 38(4):576–589.

87. Robinson MD, McCarthy DJ, Smyth GK. edgeR: a Bioconductor package for differential expression analysis of digital gene expression data. Bioinformatics 2010; 26(1):139–140.

88. Da Wei Huang BTSRAL. Systematic and integrative analysis of large gene lists using DAVID bioinformatics resources. Nature Protocols 2008; 4:44–57.

89. Subramanian A, Tamayo P, Mootha VK, Mukherjee S, Ebert BL, Gillette MA et al. Gene set enrichment analysis: a knowledge-based approach for interpreting genome-wide expression profiles. Proceedings of the National Academy of Sciences of the United States of America 2005; 102(43):15545–15550.

90. Benjamini Yoav YD. The control of the false discovery rate in multiple testing under dependency. Annals of Statistics 2001; 29:1165–1188.

91. Byers B, Cord B, Nguyen HN, Schule B, Fenno L, Lee PC et al. SNCA triplication Parkinson’s patient’s iPSC-derived DA neurons accumulate alpha-synuclein and are susceptible to oxidative stress. PloS one 2011; 6(11):e26159.

92. Wakeman DR, Hiller BM, Marmion DJ, McMahon CW, Corbett GT, Mangan KP et al. Cryopreservation Maintains Functionality of Human iPSC Dopamine Neurons and Rescues Parkinsonian Phenotypes In Vivo. Stem cell reports 2017; 9(1):149–161.

93. Banker GA. Trophic interactions between astroglial cells and hippocampal neurons in culture. Science 1980; 209(4458):809–810.

94. Banker GA, Cowan WM. Rat hippocampal neurons in dispersed cell culture. Brain research 1977; 126(3):397–342.

95. Benson DL, Huntley GW. Building and remodeling synapses. Hippocampus 2012; 22(5):954–968.

96. Sudhof TC. Towards an Understanding of Synapse Formation. Neuron 2018; 100(2):276–293.

97. Yoshii A, Constantine-Paton M. BDNF induces transport of PSD-95 to dendrites through PI3K-AKT signaling after NMDA receptor activation. Nat Neurosci 2007; 10(6):702–711.

98. Mahoney RE, Azpurua J, Eaton BA. Insulin signaling controls neurotransmission via the 4eBP-dependent modification of the exocytotic machinery. Elife 2016; 5.

99. McLaughlin CN, Broihier HT. Keeping Neurons Young and Foxy: FoxOs Promote Neuronal Plasticity. Trends Genet 2018; 34(1):65–78.

100. Al-Mubarak B, Soriano FX, Hardingham GE. Synaptic NMDAR activity suppresses FOXO1 expression via a cis-acting FOXO binding site: FOXO1 is a FOXO target gene. Channels (Austin) 2009; 3(4):233–238.

101. Bagetta V, Ghiglieri V, Sgobio C, Calabresi P, Picconi B. Synaptic dysfunction in Parkinson’s disease. Biochem Soc Trans 2010; 38(2):493–497.

102. Picconi B, Piccoli G, Calabresi P. Synaptic dysfunction in Parkinson’s disease. Adv Exp Med Biol 2012; 970:553–572.

