## Supplementary for "Reduced synaptic activity and dysregulated extracellular matrix pathways are common phenotypes in midbrain neurons derived from sporadic and mutation-associated Parkinson’s disease patients"

**Supplementary figures and legends**

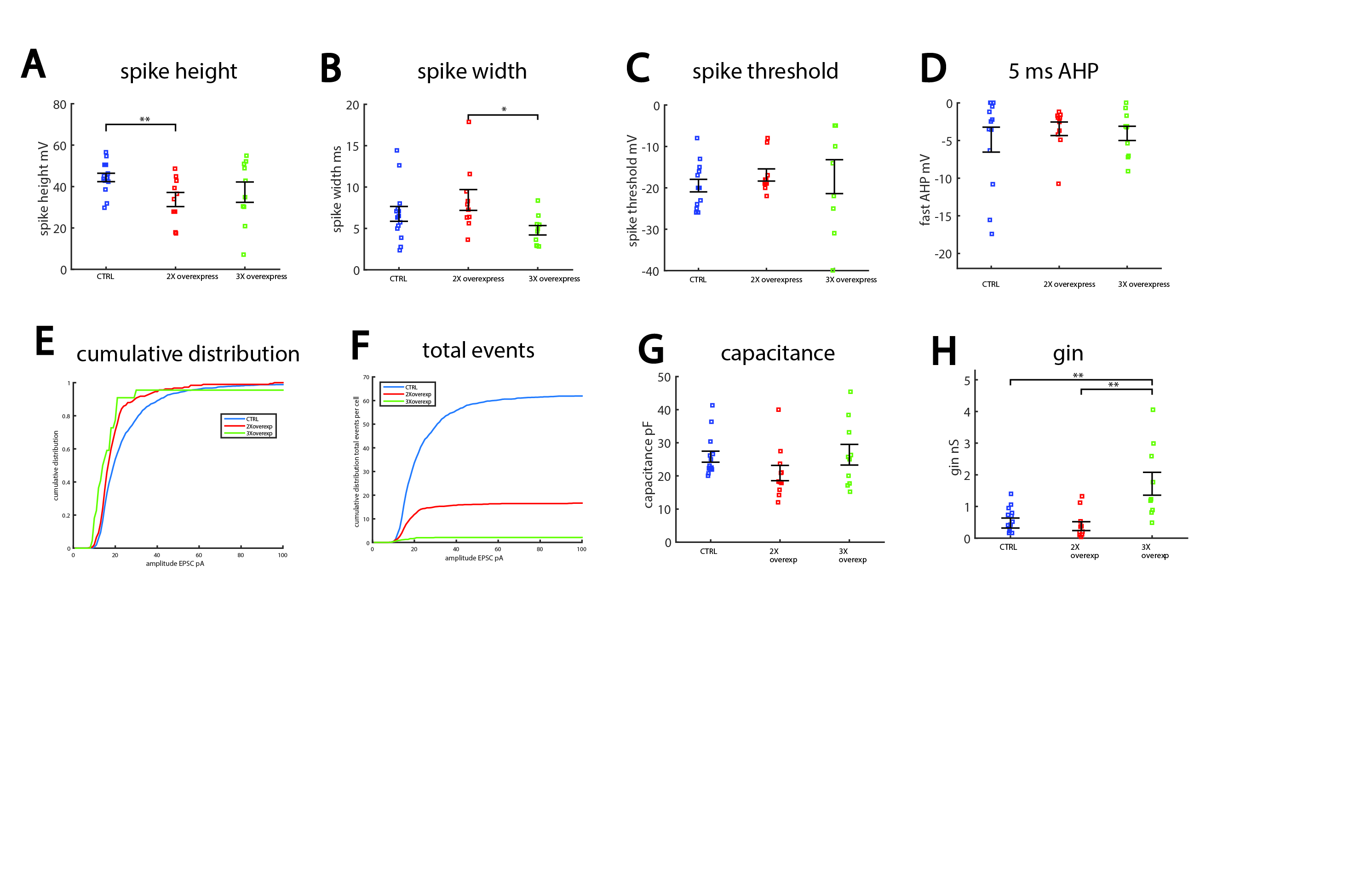

**Supplementary Figure 1.** A. Spike height is decreased in dopaminergic neurons derived from patients with a double copy of the SNCA gene compared to controls. B. Spike width is broadened in dopaminergic neurons derived from patients with a double copy (2X) of the SNCA gene compared to patients with a triple copy (3X) of the SNCA gene. C-D. No changes in the threshold for evoking an action potential and in the fast after-hyperpolarization (AHP) were observed between the 2X and 3X patient’s neurons and the controls. E. The cumulative distribution of the amplitude of synaptic currents shows that dopaminergic neurons derived from 2X and 3X patients have significantly smaller amplitude of synaptic curretns than control dopaminergic neurons. F. The total number of events per recorded cell is drastically reduced in dopaminergic neurons derived from 2X and even more in the 3X patients compared to controls. G. The capacitance is reduced (but not significantly) in the neurons derived from the 2X patient. H. The input conductance is increased in dopaminergic neurons derived from the 3X patients compared to controls and 2X patients. In this figure asterisks represent statistical significance by the following code: * p value<0.05, **p value<0.01, ***p<0.001, ****p<0.0001. Error bars represent standard error in this figure.

**Supplementary File 2.** A table summarizing all the GO terms and KEGGS significantly affected pathways in all the comparisons presented throughout the study.

**Supplementary File 3.** A complete list of all the differentially expressed genes (DEGs) in the DA neurons derived from the patient with the α-synuclein triplication.

**Supplementary File 4.** A table summarizing all the MSigDB significantly affected pathways in all the comparisons presented throughout the study.

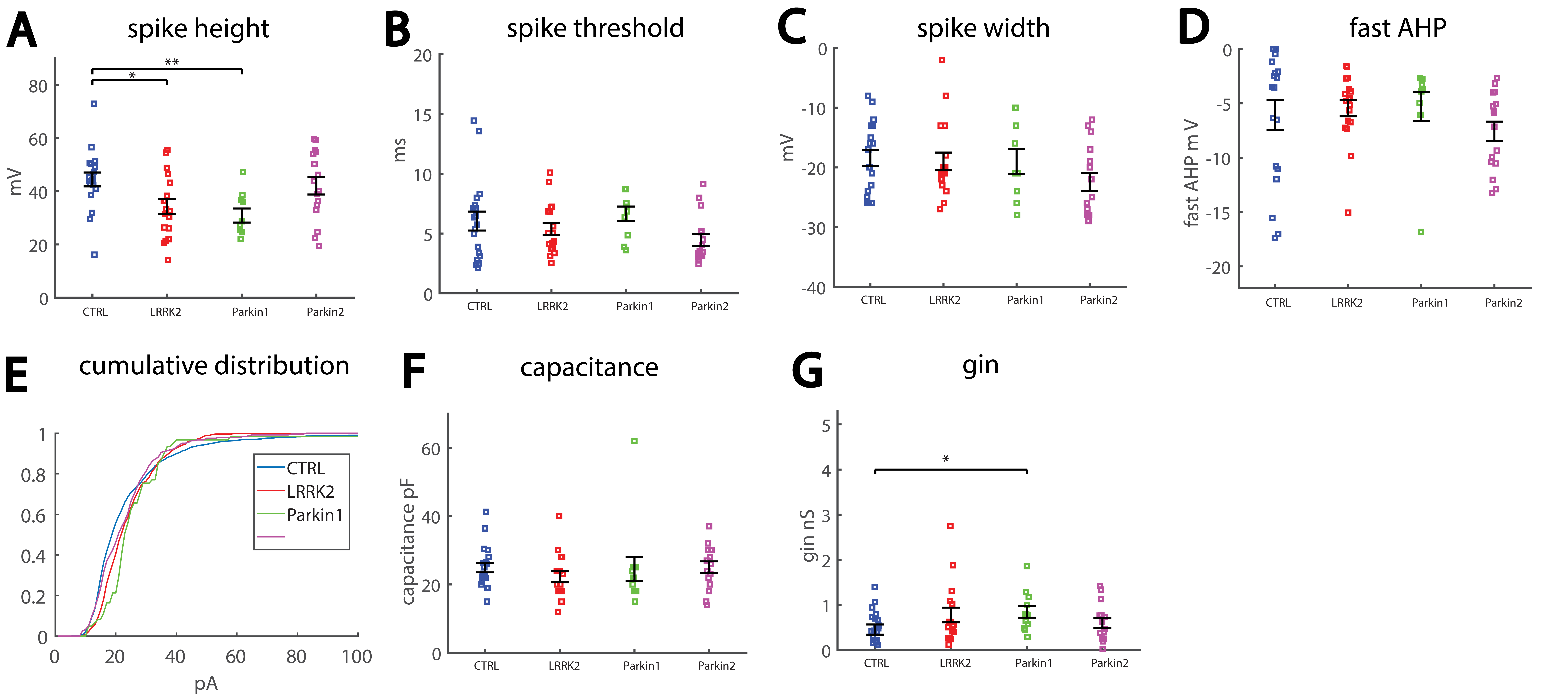

**Supplementary Figure 5.** A. Spike height was significantly smaller in dopaminergic neurons derived from the patient with the first Parkin mutation and the LRRK2 mutation, but not in the second patient with the second Parkin mutation compared to controls. B. Spike threshold was not signifcantly different between the lines. C. Spike width was unchanged between the lines. D. There was no change in the fast after-hyperpolarization (AHP) observed between the patient’s neurons and the controls. E. The cumulative distribution of the amplitude of EPSCs showing that there is no change to the distribution of amplitudes of EPSCs between control dopaminergic neurons and the ones derived from the patient with mutations in the LRRK2 and Parkin genes. F. No change was observed in the capacitance between neurons derived from healthy controls and neurons derived from the Parkin and LRRK2 mutations carrying patients. G. A significant increase was observed in the input conductance of dopaminergic neurons derived from one of the first patient with the Parkin mutation. In this figure asterisks represent statistical significance by the following code: * p value<0.05, **p value<0.01, ***p<0.001, ****p<0.0001. Error bars represent standard error in this figure.

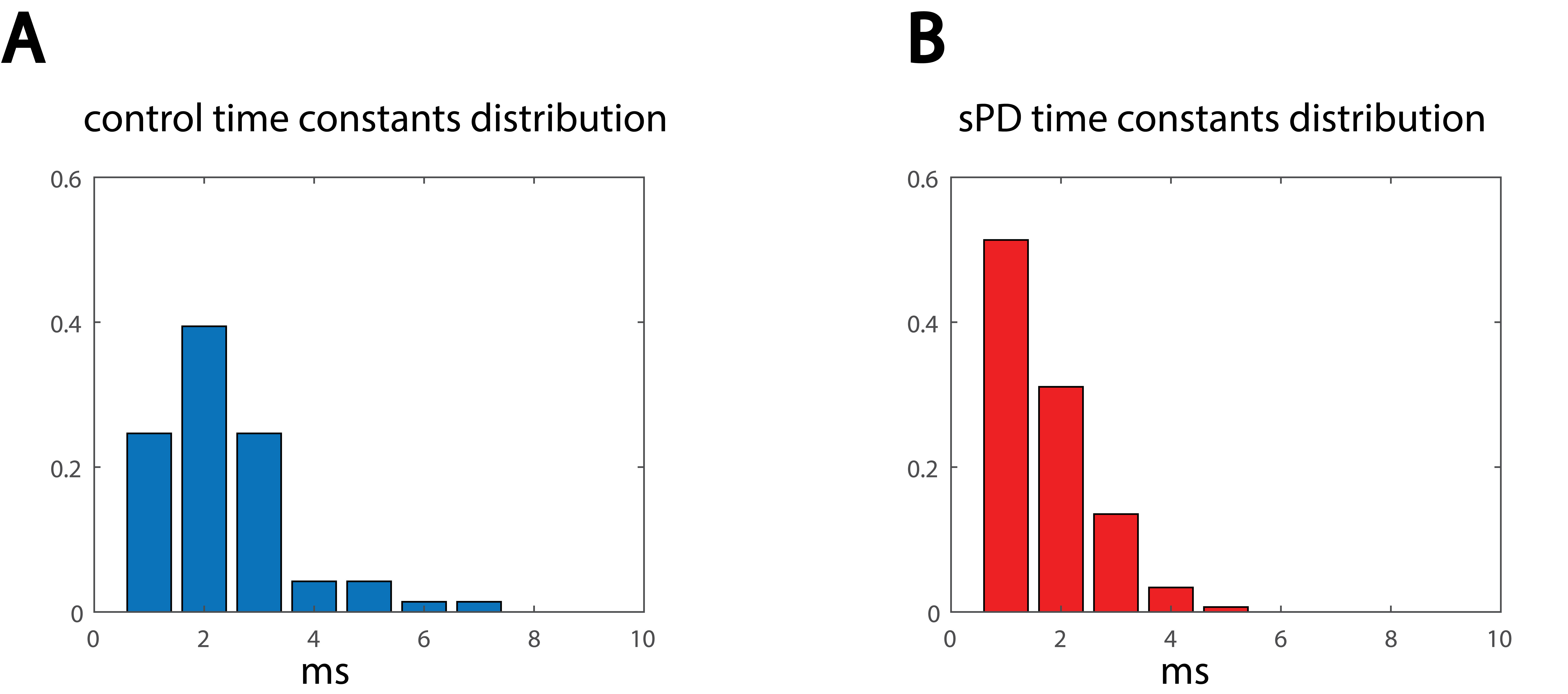

**Supplementary Figure 6.** The distribution of the decay time constants in control and sPD neurons indicates shorter decay time in sPD neurons. A. The distribution in control neurons. B. The distribution in sPD neurons.

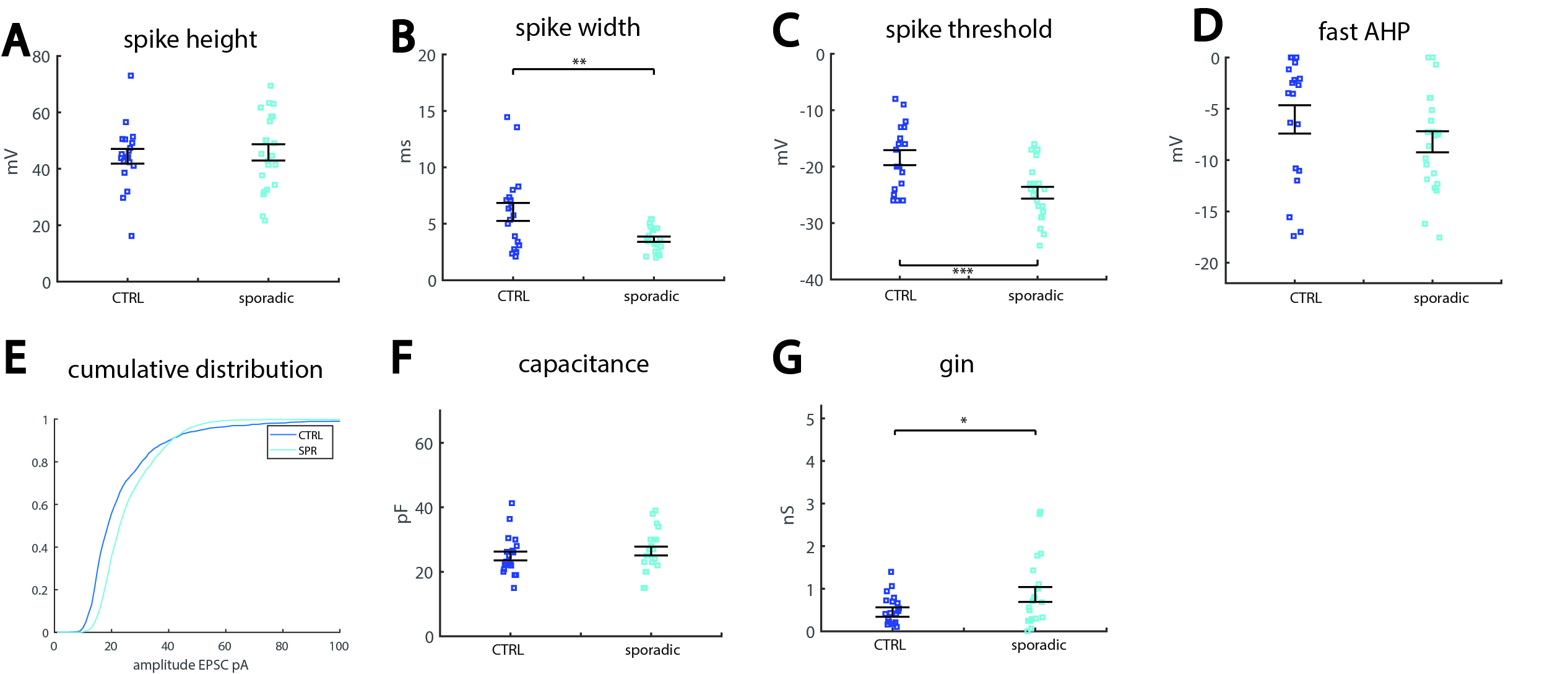

**Supplementary Figure 7** A. No significant changes were observed in the spike height between control and sPD patient. B. Spike width was significantly narrower in neurons derived from the sPD patient compared to those derived from the healthy control. C. Spike threshold was significantly less depolarized in the neurons derived from the sPD patient compared to the healthy control. D. There was no change in the fast after-hyperpolarization (AHP) observed between the patient’s neurons and the controls. E. The cumulative distribution of the amplitude of synaptic currents shows that there is no change in the distribution of amplitudes of synaptic currents between control and sPD dopaminergic neurons. F. No significant change was observed in the capacitance of the control and sPD neurons. G. A significant increase is observed in the input conductance of sPD dopaminergic neurons and controls. In this figure asterisks represent statistical significance by the following code: * p value<0.05, **p value<0.01, ***p<0.001, ****p<0.0001. Error bars represent standard error in this figure.

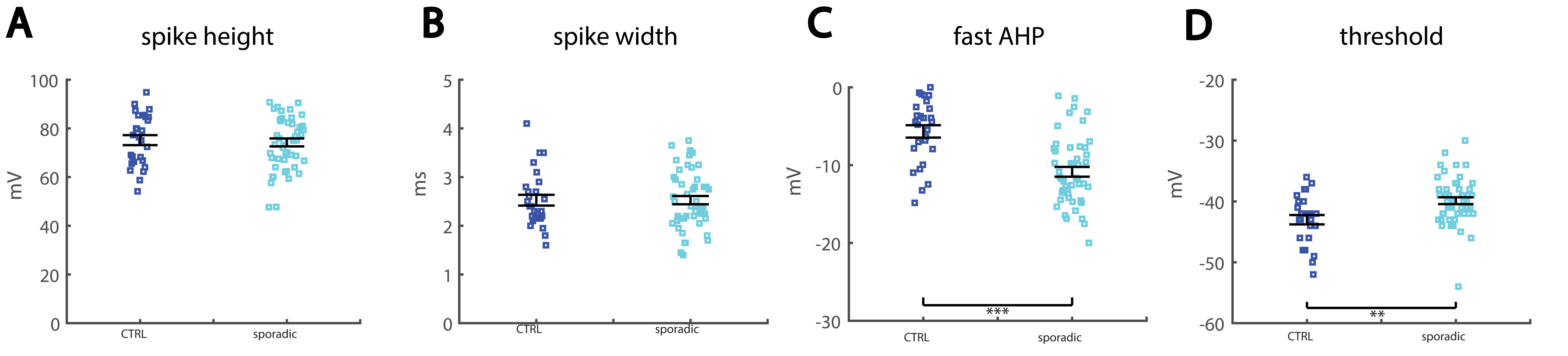

**Supplementary Figure 8.** A. No significant changes were observed in the spike height between control and sPD patient. B. No significant changes were observed in the spike width between control and sPD patient. C. An increased amplitude of the fast AHP was observed in the neurons derived from the sPD patient compared to the controls. D. A more depolarized threshold was observed in neurons derived from the sPD patient compared to the controls. In this figure asterisks represent statistical significance by the following code: * p value<0.05, **p value<0.01, ***p<0.001, ****p<0.0001. Error bars represent standard error in this figure.

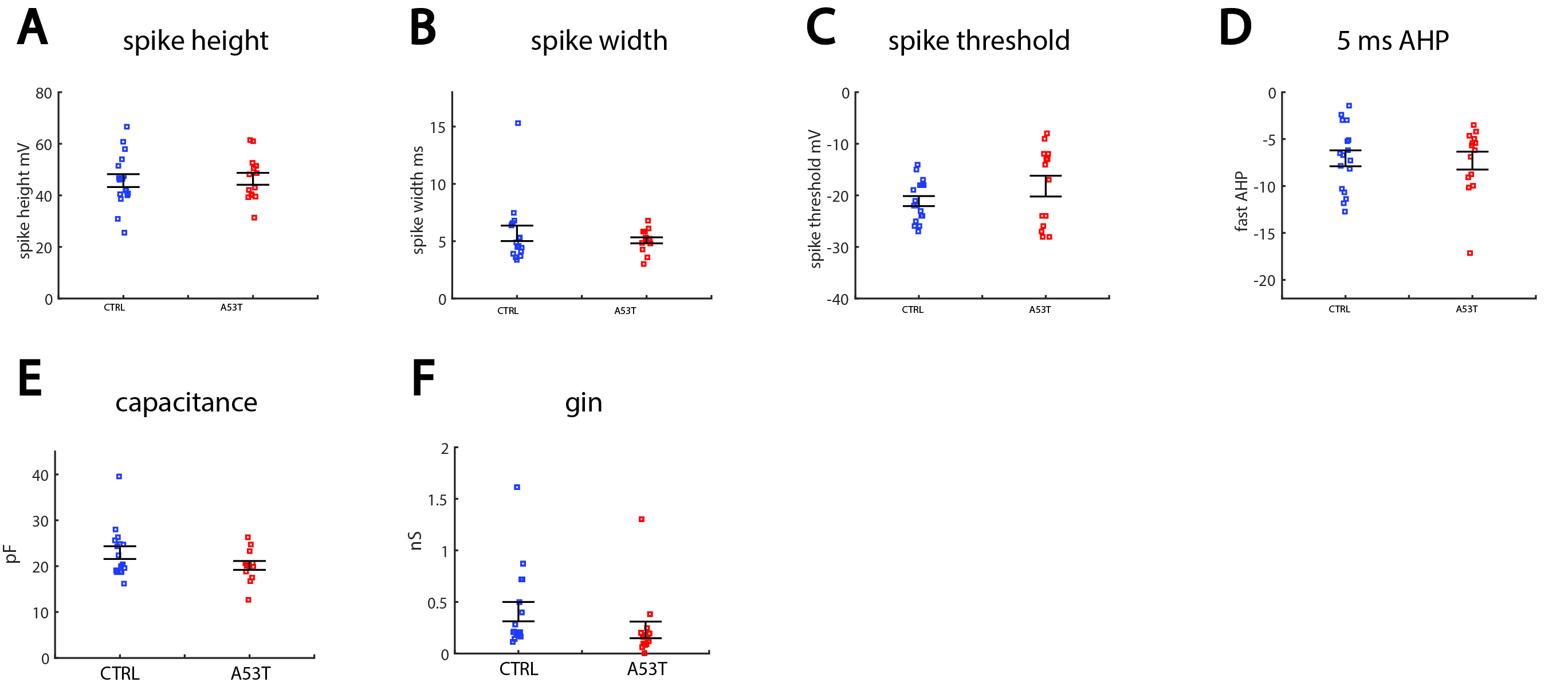

**Supplementary Figure 9.** A-F. No significant changes were observed in the spike amplitude (A), spike width (B), spike threshold (C) , the fast AHP (D), the capacitance (E) and the input conductnace (F) in the DA neurons with the A53T mutation compared to the control DA neurons.

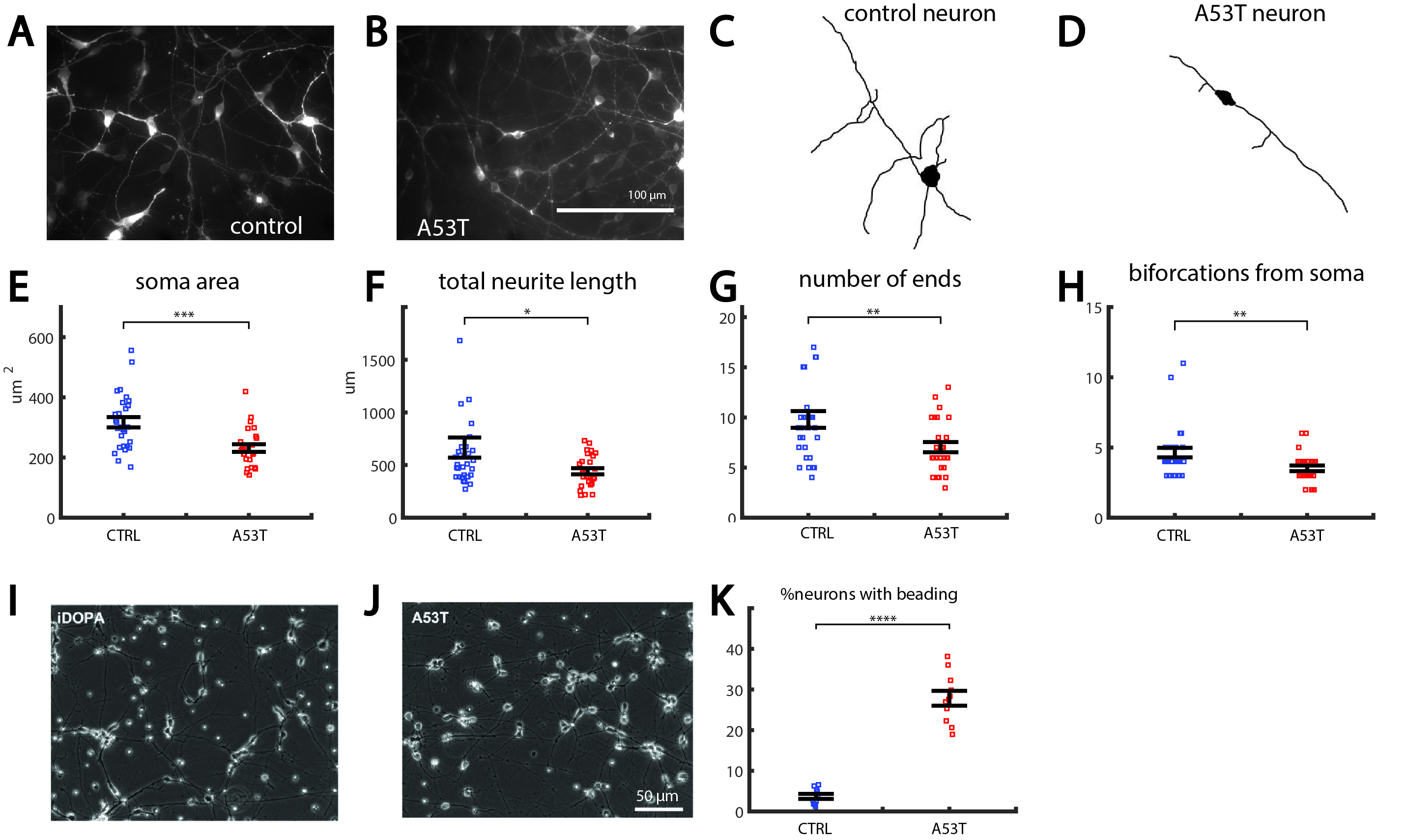

**Supplementary Figure 10.** A53T DA neurons are smaller and less arborized than control neurons. A. A representative image of a control dopaminergic neuronal culture. B. A representative image of an A53T dopaminergic neuronal culture. C. A representative traced control DA neuron. D. A representative traced A53T DA neuron. E. The soma area is reduced in A53T neurons. F. The total neurite length is reduced in A53T neurons. G. The total number of neurite ends is reduced in A53T neurons, indicating a less arborized neurite tree. H. The total number of neurites emerging out of the soma is reduced in A53T neurons. I-J. Bright field images of representative beading in A53T cultures. K. The total number of neurites with beading is significantly larger in A53T cultures. In this figure asterisks represent statistical significance by the following code: * p value<0.05, **p value<0.01, ***p<0.001, ****p<0.0001. Error bars represent standard error in this figure.
